# Charting Cervical Spinal Cord Morphometry Across the Lifespan

**DOI:** 10.64898/2026.06.03.729823

**Authors:** Kurt G. Schilling, Michael E. Kim, Matthew Amandola, Chenyu Gao, Karthik Ramadass, Praitayini Kanakaraj, Sam Bogdanov, Gaurav Rudravaram, Nancy R. Newlin, Derek Archer, Timothy J. Hohman, Angela L. Jefferson, Victoria L. Morgan, Alexandra Roche, Dario J. Englot, Murat Bilgel, Lori L. Beason Held, Luigi Ferrucci, Laurie Cutting, Laura A. Barquero, Micah A. D’archangel, Tin Q. Nguyen, Kathryn L. Humphreys, Yanbin Niu, Sophia Vinci-Booher, Carissa J. Cascio, The HABS-HD Study Team, Alzheimer’s Disease Neuroimaging Initiative, The BIOCARD Study Team, Zhiyuan Li, Daniel Moyer, Simon N. Vandekar, Panpan Zhang, Samuelle St-Onge, Sandrine Bédard, Jan Valošek, Benjamin De Leener, Julien Cohen-Adad, John C. Gore, Seth Smith, Bennett A. Landman

## Abstract

Spinal cord morphometry provides essential biomarkers of neurological health, but clinical interpretations are confounded by inter-subject variability and a lack of normative references across the full human lifespan. We address this gap by generating the first comprehensive lifespan charts for cervical spinal cord morphometry. From 78,269 T1-weighted brain MRI sessions, we identified a final cross-sectional cohort of 24,251 healthy or cognitively normal controls from 33 datasets (ages 0-100 years) with cervical cord coverage. To overcome contrast variability, we employed a state-of-the-art contrast-agnostic deep learning segmentation method, extracting cross-sectional area (CSA), anteroposterior (AP) and right–left (RL/transverse), and shape indices (AP/RL ratio, eccentricity, and solidity) from C1 to C7. Normative trajectories were modeled using Generalized Additive Models for Location, Scale, and Shape (GAMLSS). The resulting charts reveal distinct non-linear lifespan changes: rapid growth through childhood and adolescence, peak maturation occurring in early-to-mid adulthood (e.g., mid-30s for CSA), followed by gradual decreases. Significant regional variations along the cervical cord and consistent sex differences (males > females for size metrics) were quantified. Spinal cord trajectories showed strong temporal coupling with brain white matter and brainstem volumes, suggesting integrated CNS development and aging. These lifespan charts provide a robust normative framework, enabling age- and sex-specific centile scoring of individual spinal cord morphometry. This resource offers a critical tool for differentiating typical variation from pathological changes, enhancing the clinical utility of spinal cord MRI in studies of development and neurodegeneration.

## Introduction

Spinal cord morphometry provides quantitative measures reflecting macrostructural integrity that are directly relevant to neuroscience and clinical care [1–4]. Cross-sectional area (CSA), anteroposterior (AP) diameter, and right–left (RL; transverse) diameter quantify cord size and are commonly used to characterize growth and atrophy. The AP/RL ratio (sometimes called “compression” ratio) describes flattening, eccentricity describes deviation from circularity, and solidity reflects contour indentation; these shape measures may capture focal deformation not reflected by cord size alone [4–6]. These measures are altered across neurological conditions - including multiple sclerosis [7, 8], amyotrophic lateral sclerosis [9–11], spinal cord injury [12], and degenerative cervical myelopathy [5, 6] - and may support disease monitoring and prognosis. In multiple sclerosis, spinal cord atrophy is detectable from the earliest stages and correlates with motor disability [13–15]; in degenerative cervical myelopathy, automated morphometric profiling complements radiological assessment of stenosis and helps standardize detection of compressed cord [5, 6, 16]. To interpret these measurements and disentangle pathology from natural inter-subject variation, a lifespan reference for normative values is needed.

Despite this clinical potential, broad adoption is limited by measurement variability that complicates subject-specific interpretation and reduces statistical power in group studies. A century of reports - from post-mortem measurements [17, 18] to modern MRI [19–21] - has yielded disparate “normative” values for spinal cord diameters and area (for a review, see [3]), with several factors driving this inconsistency. First, biological sources such as age, sex, and individual anatomical differences produce between-person dispersion [1, 22–25]. Second, methodological sources introduce bias across studies, including scanner vendor, acquisition contrast/protocol, and analysis software assumptions about cord boundaries [26–28]. Finally, variability arises from spatial localization, encompassing both the precision of identifying anatomical landmarks and the inherent neuroanatomical variance between vertebral levels and functional spinal cord segments [1]. Together these effects hinder cross-site aggregation, meta-analysis, and reliable longitudinal tracking.

Normative modeling addresses these challenges by adapting the logic of pediatric growth charts [29, 30] to neuroimaging. Recent brain-wide “charts” provide age- and sex-stratified trajectories and variance across the lifespan [31, 32]. Crucially, such models estimate the full population-level distribution of a given biological measure - not just the mean - so individual measurements can be expressed as centile scores relative to age-appropriate expectations, enabling standardized detection of atypical development or atrophy [31].

The main barrier to creating comprehensive spinal cord charts has been the lack of large-scale, dedicated imaging datasets spanning the full human lifespan. While valuable prior work laid the foundation by creating normative databases for specific populations, such as the PAM50 spinal cord template for adults [20, 21, 33], and by exploring normalization strategies to reduce inter-subject variability in adult and pediatric cohorts [1, 5, 6, 23, 34], these efforts were often limited by narrow age ranges, modest sample sizes, a focus on single vertebral levels, or simplified models unable to capture age-varying variance. Here, we overcome these constraints by leveraging numerous large-scale, population-based brain neuroimaging datasets whose imaging protocols included cervical cord coverage. The feasibility and reliability of using such data for upper cervical cord morphometry are well-established [35, 36], presenting an exemplary opportunity to utilize existing data resources. However, aggregating data across these diverse cohorts also introduces significant methodological challenges, primarily MRI contrast variability (across sites or contrasts). To address this, we employ a recently developed, state-of-the-art, contrast-agnostic segmentation method [2, 37], which provides consistent morphometric estimates across different scanners, sites, and protocols.

Our goal is to create vertebral-level, sex-stratified normative charts of cervical spinal cord morphometry across the lifespan. From an initial pool of 78,269 T1-weighted MRI sessions, we assembled a cross-sectional cohort of 24,251 healthy or cognitively normal controls from 33 neuroimaging datasets spanning ages 0-100 years. we (i) model the non-linear trajectories of spinal cord growth, maturation, and atrophy; (ii) identify key lifespan milestones (e.g. age of peak CSA and diameters); (iii) quantify annualized (cross-sectional) rates of change for clinical benchmarking across life stages; (iv) characterize population variability; (v) delineate sex-specific trajectories; and (vi) relate cord morphometry to brain volumes across development and aging.

## Materials and Methods

This study involved three main stages: (i) aggregation and curation of a large-scale, multi-site neuroimaging dataset; (ii) automated processing of all images to extract spinal cord and brain morphometric features; and (iii) normative modeling of these features to generate lifespan charts.

### Study Cohorts

This work builds upon prior large-scale data aggregation efforts that collated imaging from publicly available and locally acquired neuroimaging studies [38]. The imported pool comprised 78,269 T1-weighted MRI sessions from 48,130 dataset-specific participants spanning ages 0-100 years.

We first retained sessions from participants designated by their source cohorts as healthy or cognitively normal controls, yielding 41,784 sessions from 26,697 participants. Visual quality assurance (see below) retained 37,888 sessions from 24,437 participants. After removing individuals with missing age/sex, we created a final cross-sectional modeling dataset by randomly selecting a single scan from each participant with multiple time points, a strategy consistent with large-scale charting initiatives [31]. This yielded a final cohort of 24,251 participants from 33 datasets (**Figure 1A, 1B**). Additional dataset information, sample sizes, and age ranges are provided in **Supplementary Table 1**. T1w imaging parameters are given in **Supplementary Table 2**.

**Figure 1.**
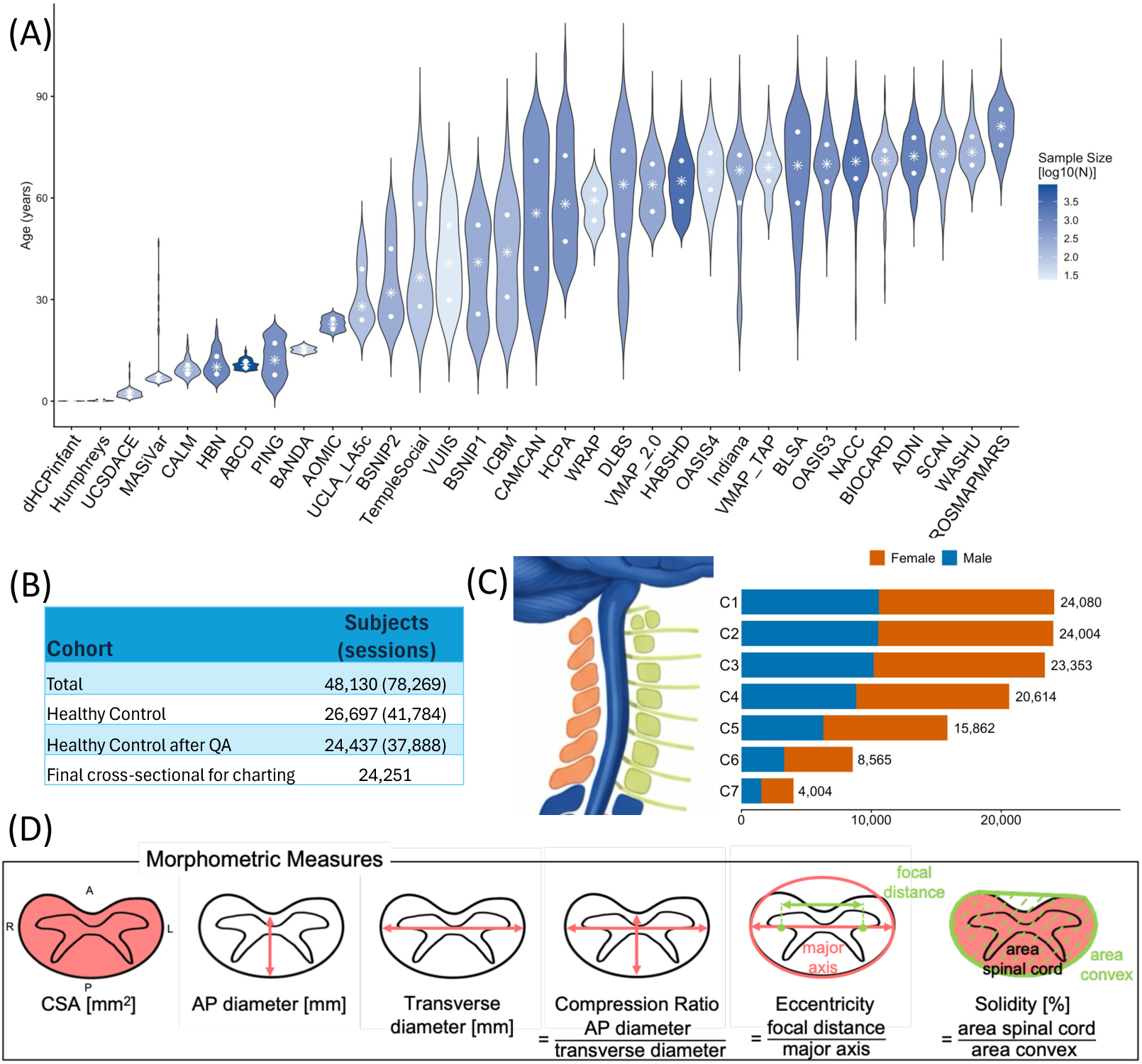
Overview of study cohorts and extracted spinal cord morphometry. Age distributions are shown across 33 contributing datasets for the final 24,251 participants used to create the spinal cord charts (A). Of 48,130 dataset-specific participants (78,269 sessions), 26,697 participants (41,784 sessions) were designated healthy or cognitively normal controls, 24,437 participants (37,888 sessions) remained after scan-level QA, and selection of one eligible scan per participant yielded 24,251 participants from 33 datasets for charting (B). The number of participants contributing to the CSA models varied by vertebral level (C). Six morphometric features were extracted at each available level: cross-sectional area (CSA), anteroposterior (AP) diameter, right-left (RL) diameter, AP/RL ratio, eccentricity, and solidity (D; cord illustration recreated from [21]).

All vertebral labeling and segmentation outputs underwent visual quality control (**Supplementary Figure 1**); labeling failures were excluded and segmentation failures at the affected level only (i.e., failures were more common inferiorly), so a measurement contributed only when the corresponding level passed QA. Because cervical coverage varied across brain T1-weighted acquisitions, the final sample sizes range from C1=24,080 to C7=4,004 samples. (**Figure 1C**); their age-by-sex distributions are shown in **Supplementary Figure 2**. We note that levels C6/C7 have sparse data throughout young adulthood.

### Image processing

#### Spinal cord morphometry

All T1-weighted images were processed with the Spinal Cord Toolbox (SCT, v7.1) [39]. To handle variability in image contrast across the aggregated datasets, we used a state-of-the-art, contrast-agnostic deep learning model to automatically segment the spinal cord from all T1-weighted images (*sct_deepseg spinalcord*) [2]. This method offers a key advantage critical for this large-scale study: it provides robust and consistent segmentation performance regardless of the input image contrast (i.e., differences in acquisition parameters across sites). Vertebral levels were extracted using a tool for automatic instance segmentation of all vertebrae (*sct_deepseg totalspineseg*), that is also robust to various MRI contrasts, acquisition orientations, and resolutions. Following SCT convention, C1 and C2 were retained as separate projected vertebral regions; because no intervertebral disc separates the atlas and axis, their boundary was estimated from upper-cervical vertebral landmarks [21, 39].

Following segmentation, three measures of cord size (CSA and AP- and RL-diameters) and three measures of cross-sectional shape (AP/RL ratio, eccentricity, and solidity) were extracted at each vertebral level in native space using *sct_process_segmentation* (**Figure 1D**):

- Cross-Sectional Area (CSA) (mm²): The total area of the spinal cord in the axial plane (on a centerline-orthogonal axial slice), calculated from the binary segmentation by summing the voxel-wise probabilities. CSA is a primary indicator of spinal cord atrophy.
- Anterior-Posterior (AP) Diameter (mm): The maximum diameter of the spinal cord along the anterior-posterior axis. Calculated as the AP axis of a fitted ellipse.
- Right-Left (RL) / Transverse Diameter (mm): The maximum diameter of the spinal cord along the right-left axis, also referred to as the transverse diameter. Calculated as the RL axis of a fitted ellipse.
- AP/RL Ratio (a.u.): The ratio between the AP and RL diameters, which provides a measure of cord flattening (lower value indicates greater flattening). This had previously been referred to as “compression” ratio in literature and within SCT [21], but is called AP/RL ratio here as its name may inappropriately suggest spinal cord compression.
- Eccentricity (a.u.): A measure of the spinal cord’s deviation from a perfect circle, calculated from an ellipse with the same second-order moments as the cord segmentation (range 0–1; values nearer 0 indicate a more circular cross-section).
- Solidity: The ratio of the CSA to the area of its convex hull, which quantifies the convexity of the cord shape and is sensitive to local indentations (lower values indicate greater indentation/non-convexity)

#### Brain volumetrics

To investigate the relationship between brain and spinal cord morphometry, we processed all T1-weighted images using FreeSurfer (*recon-all*, v7.2.0) [40]. FreeSurfer segmentations were visually quality controlled and failed or unusable outputs were excluded as previously described [38]; dataset-level terms in the normative models were used to account for scanner- and acquisition-related variation in the derived volumes [41]. For comparative analysis with spinal cord metrics, we extracted the following primary brain volumes: Total Brain Volume, Total Gray Matter Volume, Total White Matter Volume, Thalamus Volume (right + left), and Brainstem Volume.

To investigate cross-sectional brain–cord relationships, we compared median lifespan trajectories and calculated Pearson correlations between observed brain and spinal cord measures within each lifespan stage. Correlations were evaluated across C1–C7 and for all cord measures; coefficients were interpreted descriptively without multiple-comparison testing.

#### Normative modeling and lifespan charting

We modeled age-dependent normative trajectories using Generalized Additive Models for Location, Scale, and Shape (GAMLSS) [42] (as implemented in R as the gamlss package), following the framework adopted for pediatric growth [43] and recent large-scale brain charts [31, 44]. Each outcome (per-level cord morphometric or global brain volume) was fit with the Generalized Gamma (GG) family, with the location (μ) and scale (σ) parameters expressed as smooth, non-linear functions of age and sex. Age effects were modeled with fractional polynomials (order 2 for μ; order 1 for σ), and sex entered both parameters as an additive fixed effect; no age-by-sex interaction was fitted. To account for between-study/site heterogeneity, we included dataset-level random effects in both μ and σ, yielding site-robust population estimates while retaining within-dataset information. In the GG family, the shape parameter (ν) - which governs distributional skewness - was treated as constant (i.e., not modeled as a function of covariates) in our fits. Model adequacy was assessed independently using standard GAMLSS diagnostics, including worm plots and empirical-vs-model centile checks, to verify that fitted curves reproduced observed trends.

Spinal cord measures were modeled in native units without intracranial-volume normalization so that the charts describe age- and sex-specific population variation and remain applicable to studies without brain imaging.

Outlier handling prior to fitting used two safeguards: (i) dataset- and level-specific screening for implausible values suggestive of segmentation or coverage error, and (ii) exclusion of observations exceeding ±3 SD relative to the dataset-level mean for that feature and level. Models were fit on the healthy cross-sectional cohort, separately for each vertebral level (C1– C7) and morphometric feature.

The output was a set of continuous, age- and sex-specific centile curves that constitute the spinal-cord lifespan charts, describing the population distribution via μ, σ (and constant ν). For visualization we show canonical centiles (2nd, 5th, 10th, 25th, 50th, 75th, 90th, 95th, 98th).

From the fitted median (50th-centile) curve we derive two summaries used throughout the Results: the annualized rate of change (from the first derivative; also expressed as %/year) and the age of peak (maturation; maximum of the fitted median curve). Because these rates are derived from cross-sectional data, they represent population-level age associations and should not be interpreted as within-person longitudinal change.

For interpretability, rates are summarized across predefined lifespan stages: Infancy (0–1 y), Early Childhood (1–6 y), Late Childhood (6–12 y), Adolescence (12–20 y), Young Adulthood (20–40 y), Middle Adulthood (40–60 y), and Late Adulthood (60–100 y).

#### Model stability and robustness

To assess sensitivity to scan selection, we refitted the models after randomly selecting one eligible scan per participant ten times. We also performed leave-one-dataset-out refits; variability in the median centile trajectories was summarized using the coefficient of variation across refits.

#### Out-of-sample alignment

While a new individual’s morphometric measure can simply be placed on our charts (as in pediatric charting), it is advantageous to also enable a dataset-level harmonization to account for possible scale/site/acquisition differences.

To demonstrate application of the spinal cord charts to a new study, we performed out-of-sample alignment of an exemplar dataset excluded from the reference-chart used for this analysis. We chose an in-house dataset (VUIIS-MS) collected to study multiple sclerosis (MS), a chronic inflammatory and neurodegenerative disorder of the central nervous system in which spinal cord involvement and atrophy are common and associated with clinical disability and disease progression [7, 13, 15, 27, 37, 45]. T1-weighted images were processed using the same spinal cord segmentation, vertebral labeling, quality-control, and morphometric procedures used for chart construction. This cohort included 67 participants: 34 controls (age 22.7-64.1; mean age 38.6; 26 female) and 33 participants with MS (age 25.0-64.4; mean age 42.4; 29 female). We note participants with MS generally had minimal disability (mean EDSS=2) and were on disease modifying treatments.

Following previous normative charting studies [31, 38], dataset-specific offsets to the location and scale of the generalized gamma reference distribution were estimated from controls using maximum-likelihood estimation. These offsets were applied unchanged to participants with MS, after which age- and sex-specific centiles were calculated for every participant. The same procedure can be applied to any dataset containing age, sex, control-group labels, and any morephometric feature.

To validate the procedure, we repeated the analysis after introducing a 5% multiplicative bias in C3 CSA. We also evaluated alignment in a larger ADNI sample of 1,157 controls and 748 participants with Alzheimer disease. The reference charts, deployable model files, alignment software, and worked examples are distributed through Zenodo (doi:10.5281/zenodo.18306654).

## Results

### Non-linear Dynamics of Spinal Cord Morphometry Across the Lifespan

We generated normative lifespan charts for cervical spinal cord morphometry by applying GAMLSS to our large-scale, cross-sectional cohort. At the representative C3 level (**Figure 2**), the raw data and fitted centiles show nonlinear trajectories across the lifespan. CSA, AP-diameters, and RL (transverse)-diameters rise rapidly through childhood and adolescence, plateau in young adulthood, and decreases in later life. Other shape metrics follow different patterns: AP/RL ratio generally decreases with age, eccentricity increases modestly, and solidity is largely stable with a slight late-life reduction.

**Figure 2.**
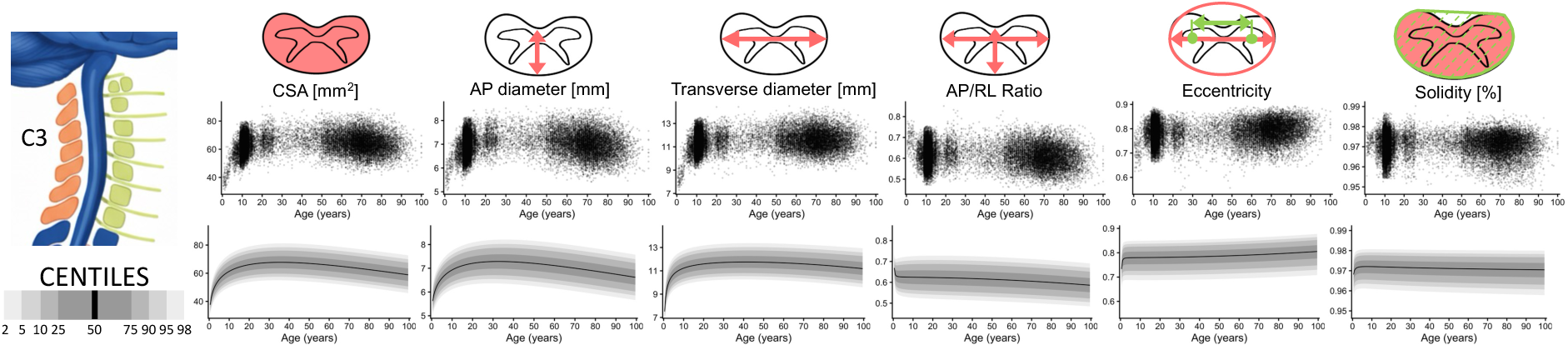
Cervical Spinal Cord Growth Charts Across the Lifespan. Lifespan trajectories for six morphometric features are shown at the representative C3 vertebral level. Individual data points from the healthy cohort are plotted, along with the estimated centile curves (2nd, 10th, 25th, 50th, 75th, 90th, 98th) derived from GAMLSS models. Measures of cord size, including Cross-Sectional Area (CSA), Anterior-Posterior (AP) Diameter, and Right-Left (RL) Diameter, follow a characteristic non-linear trajectory of rapid growth through childhood, a plateau in young adulthood, and a gradual decrease in later life. In contrast, measures of cord shape exhibit distinct lifespan dynamics: AP/RL Ratio shows a subtle decline with age, Eccentricity demonstrates a gradual increase, and Solidity remains largely stable before a slight decrease in late adulthood.

**Figure 3** extends these results across C1–C7 for CSA and AP diameter, presenting (i) the normative lifespan trajectory (median), (ii) the annualized rate of change derived from the first derivative, and (iii) population variability (σ). While the global pattern (growth → plateau → decrease) is conserved, the timing and magnitude vary by vertebral level. Parallel panels for RL diameter, AP/RL ratio, eccentricity, and solidity are provided in **Supplementary Figure 3**, and complete centile tables for all features, levels, and sexes are provided as **Supplementary Data**.

**Figure 3.**
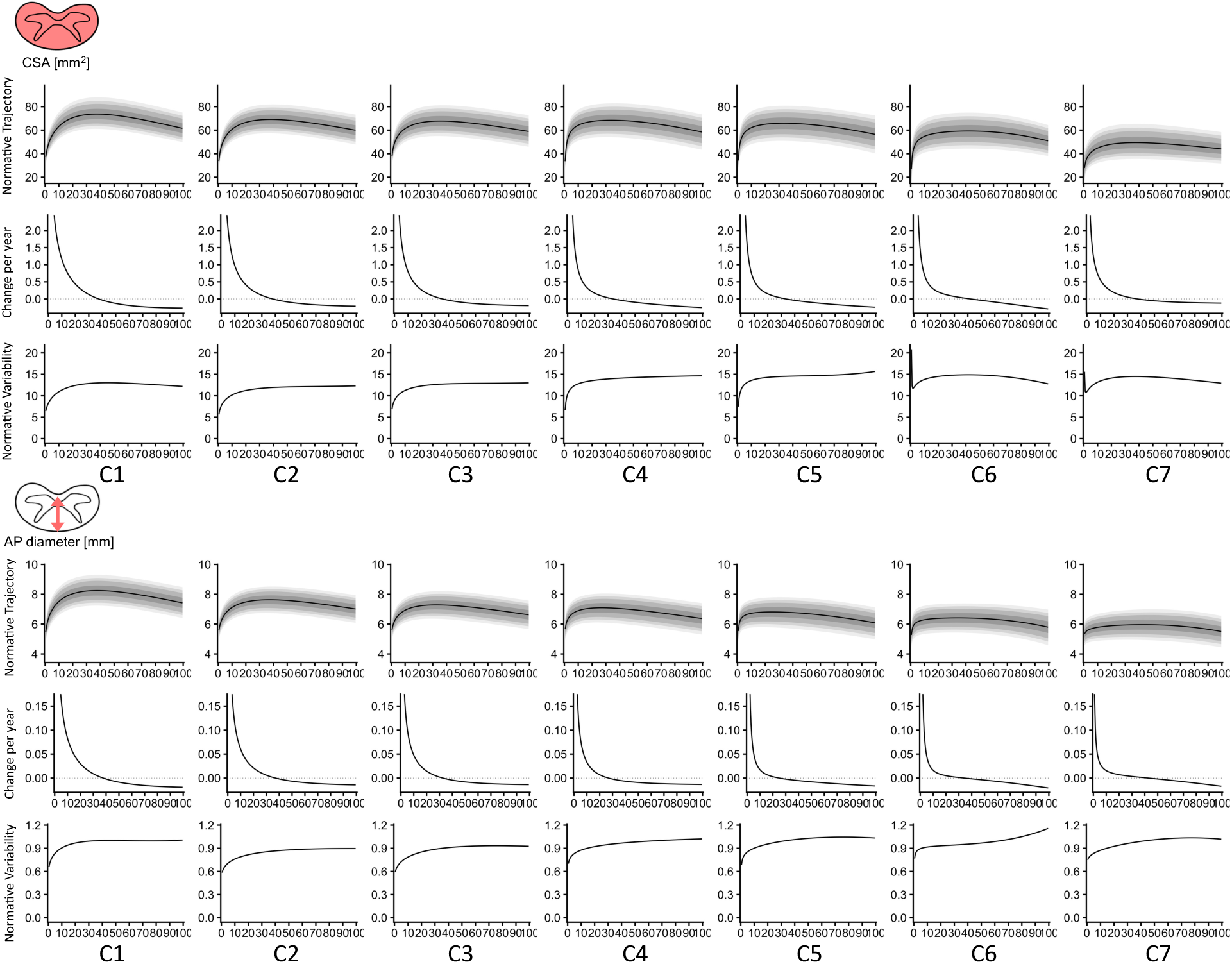
Lifespan Dynamics of Spinal Cord Morphometry Across the Vertebral Levels. Figure displays the normative trajectory (centiles), annualized rate of change (change-per-year), and normative population variability (standard deviation, σ) for Cross-Sectional Area (CSA, top) and Anterior-Posterior (AP) Diameter (bottom). Normative trajectories reveal distinct developmental and aging patterns that vary by vertebral level; annualized rate of change quantifies these dynamics, highlighting periods of rapid growth in childhood and adolescence followed by cross-sectional decreases in late adulthood; Population variability illustrates the spread of measurements changes across the lifespan.

### Regional Variations Along the Spinal Cord with Age

The normative lifespan charts reveal level-dependent patterns across C1-C7 vertebral levels (**Figure 4**). Size metrics (CSA, AP, RL) generally decrease caudally, with a prominent cervical enlargement at C4–C5 driven chiefly by greater RL (transverse) diameter, consistent with neuroanatomy [18, 21]. While the absolute magnitude of size metrics varies across levels, the overall shape of the lifespan trajectory - early growth and late-life decrease - is largely conserved along the cord. In contrast, measures of cord shape, such as AP/RL ratio and eccentricity, exhibit more complex and level-specific lifespan dynamics, with unique patterns of development and aging at different vertebral locations.

**Figure 4.**
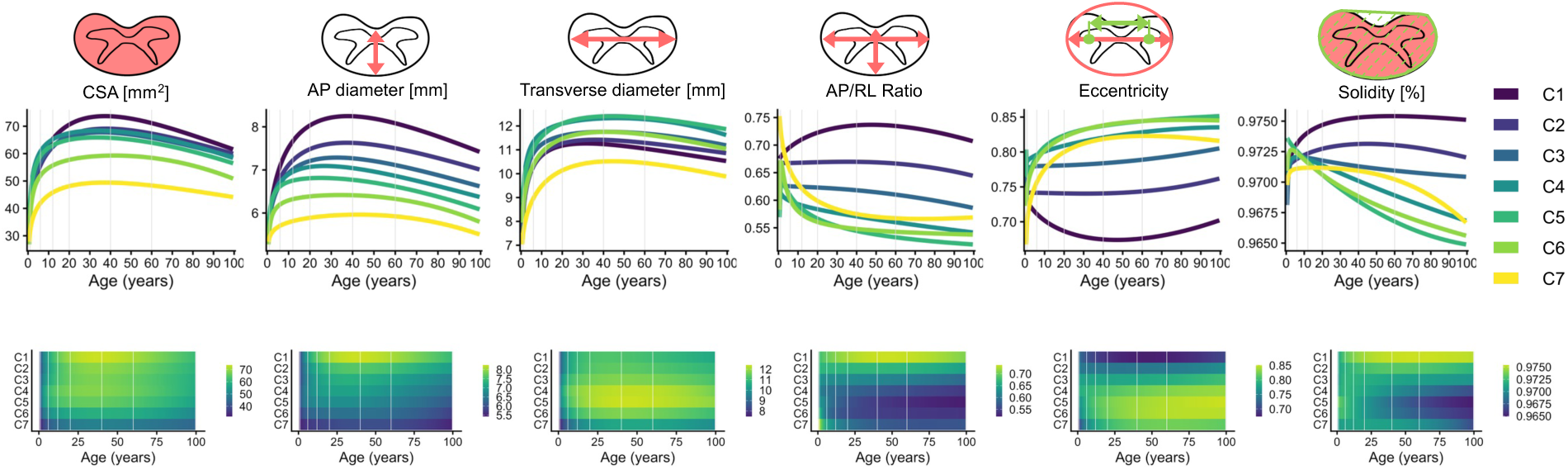
Regional variations along the spinal cord with age. Median lifespan trajectories for all six morphometric features are plotted for each cervical vertebral level (C1–C7) (top row), alongside two-dimensional heatmaps visualizing the magnitude of each feature across both age and vertebral level (bottom row). While overall trends are shared across levels, the timing and magnitude of change are level-dependent, and shape metrics (AP/RL ratio, eccentricity, solidity) display more heterogeneous, region-specific dynamics.

### Lifespan Milestones in Spinal Cord Growth and Aging

The normative lifespan charts allow for the identification of key developmental milestones, such as the age at which spinal cord morphometric measures reach their peak (Figure 5), which can be interpreted as the age of maturation [23]. CSA peaked between 33 and 42 years across cervical levels (35 years at C3), refining a prior estimate of approximately 45 years from a smaller single-site adult cohort (N=129) [23]. AP diameter generally peaked earlier (25–43 years) than RL diameter (31–47 years), particularly at C3–C7. These peaks provide quantitative benchmarks for the transition from growth to age-related decline.

**Figure 5.**
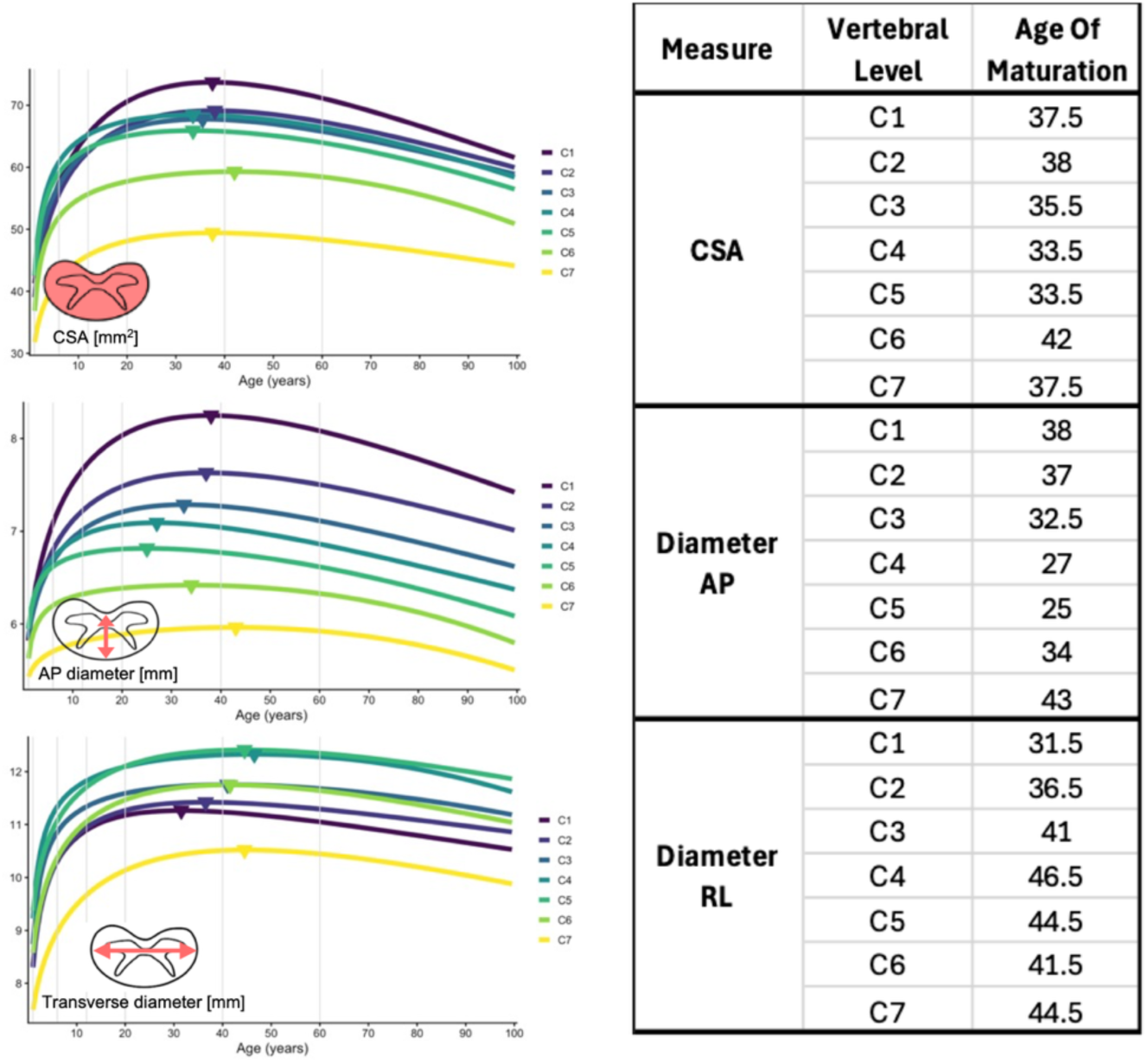
Lifespan milestones in spinal cord growth and maturation. The median lifespan trajectories for Cross-Sectional Area (CSA), Anterior-Posterior (AP) Diameter, and Right-Left (RL) Diameter are shown for each cervical vertebral level. The peak of each curve, interpreted as the age of maturation, is indicated by a marker. The peak of each curve, interpreted as the age of maturation, is marked and quantified in the accompanying table. CSA peaked between 33 and 42 years across levels, while AP diameter generally peaked earlier than RL diameter, particularly at C3–C7.

### Quantifying Annualized Rates of Change for Clinical Benchmarking

To establish quantitative benchmarks for clinical and research applications, we derived the annualized rates of change from the (cross-sectional) normative curves for each morphometric feature across distinct lifespan stages (**Table 1**). Change was greatest in early life: CSA increased dramatically in infancy and by 6.7–9.5%/year during early childhood; over the same period, RL diameter increased more rapidly than AP diameter at every level (3.9–5.7%/year versus 1.1–4.4%/year). A broad plateau followed, whereas in late adulthood CSA decreased by 0.23–0.37%/year; AP diameter declined by 0.17–0.22%/year, exceeding the 0.10–0.14%/year decline in RL diameter at every cervical level. Absolute rates (mm²/year and mm/year) and stage-wise results for shape metrics are provided in **Supplementary Table 3**.

**Table 1.**
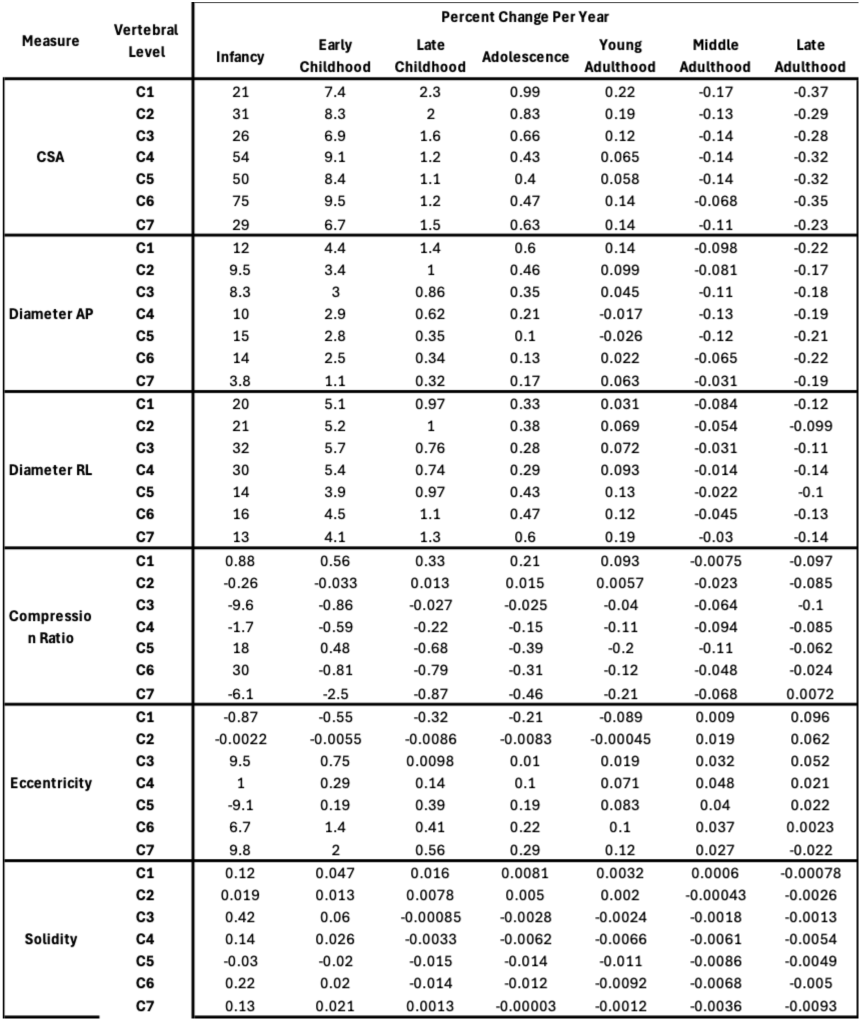
Annualized percentage rates of cross-sectional age-related change in spinal cord morphometric measures.

### Charting Population Variability Across the Lifespan

To characterize normative population variability, we calculated the coefficient of variation (CV) by lifespan stage (**Table 2**). Outside infancy, CSA CVs ranged from 8–12% at C1–C5 and were higher at C6–C7 (13–15%; possibly due to smaller sample sizes and increased susceptibility to algorithmic variance at the edge of the imaging field-of-view), while AP and RL diameter CVs remained below 10% across all levels (5.3–9.9%). CVs were largely stable across age, with modest increases at several levels in middle and late adulthood. Stage-wise variability (standard deviation and CV) for all features and levels is reported in **Supplementary Table 4**.

**Table 2.** Population Variability of Spinal Cord Size Metrics Across the Lifespan.

| Measure | Vertebral Level | Coefficient of Variation |  |  |  |  |  |  |
| --- | --- | --- | --- | --- | --- | --- | --- | --- |
|  |  | Infancy | Early Childhood | Late Childhood | Adolescence | Young Adulthood | Middle Adulthood | Late Adulthood |
| CSA | C1 | 8.6 | 8.6 | 8.6 | 8.7 | 8.7 | 8.9 | 9.4 |
|  | C2 | 8.4 | 8.4 | 8.4 | 8.4 | 8.5 | 8.8 | 9.6 |
|  | C3 | 9.1 | 9.2 | 9.2 | 9.2 | 9.3 | 9.6 | 10 |
|  | C4 | 9.9 | 9.9 | 9.9 | 10 | 10 | 11 | 12 |
|  | C5 | 11 | 11 | 11 | 11 | 11 | 11 | 12 |
|  | C6 | 39 | 13 | 13 | 13 | 13 | 13 | 13 |
|  | C7 | 28 | 15 | 15 | 15 | 15 | 15 | 15 |
| Diameter AP | C1 | 6 | 6 | 6 | 6 | 6 | 6.1 | 6.4 |
|  | C2 | 5.2 | 5.3 | 5.3 | 5.4 | 5.6 | 5.8 | 6.2 |
|  | C3 | 5.3 | 5.4 | 5.6 | 5.8 | 6.1 | 6.4 | 6.8 |
|  | C4 | 6.2 | 6.2 | 6.3 | 6.4 | 6.7 | 7 | 7.6 |
|  | C5 | 6.2 | 6.4 | 6.7 | 6.9 | 7.3 | 7.7 | 8.2 |
|  | C6 | 7.3 | 7.3 | 7.3 | 7.3 | 7.3 | 7.6 | 8.7 |
|  | C7 | 7 | 7.2 | 7.5 | 7.7 | 8.1 | 8.5 | 9 |
| Diameter RL | C1 | 7.7 | 6 | 5.7 | 5.7 | 5.6 | 5.6 | 5.6 |
|  | C2 | 5.7 | 5.7 | 5.7 | 5.7 | 5.7 | 5.8 | 6.2 |
|  | C3 | 6.4 | 6.4 | 6.4 | 6.4 | 6.4 | 6.5 | 6.8 |
|  | C4 | 6.4 | 6.4 | 6.5 | 6.6 | 6.8 | 7.1 | 7.7 |
|  | C5 | 7.1 | 7.1 | 7.1 | 7.1 | 7.1 | 7.3 | 7.9 |
|  | C6 | 8 | 8 | 8 | 8 | 8 | 8.1 | 8.6 |
|  | C7 | 18 | 9.9 | 9.6 | 9.5 | 9.5 | 9.5 | 9.5 |

### Sex Differences in Spinal Cord Morphometry

To characterize sex-specific differences, we modeled lifespan trajectories for males and females by including sex as a fixed-effect for both location and scale (Figure 6). Males generally had larger CSA and diameters, although the magnitude varied by feature and level, whereas shape metrics showed smaller sex effects. These findings motivate sex-specific centiles; age-by-sex interactions were not evaluated.

**Figure 6.**
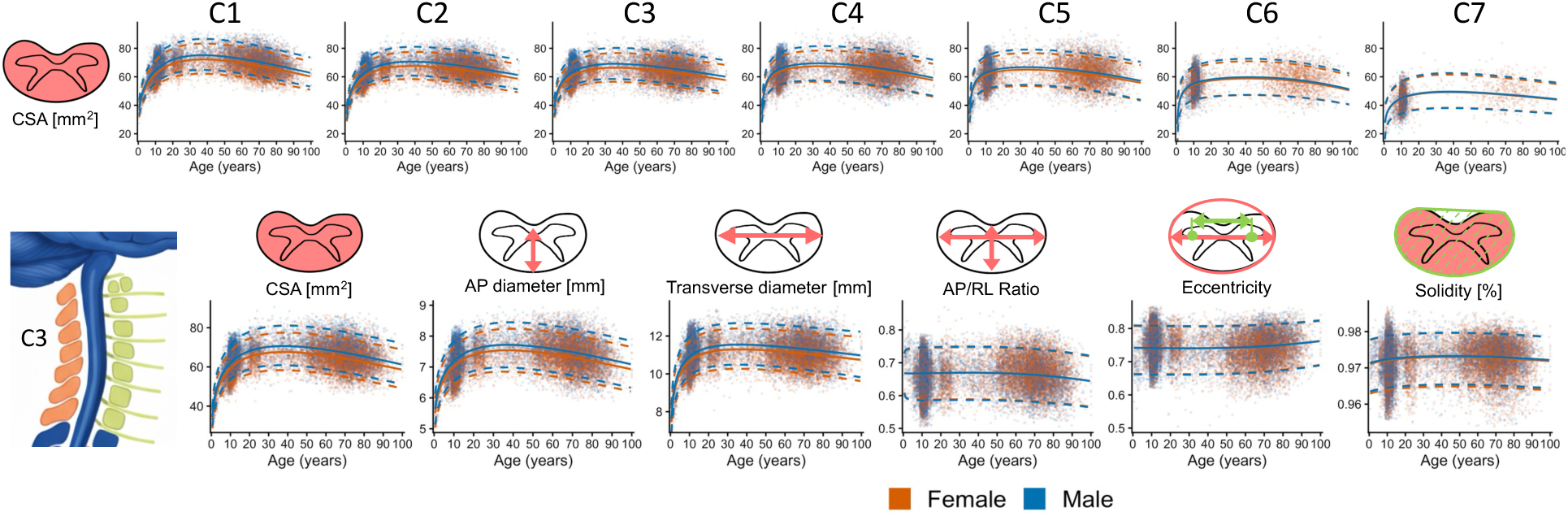
Sex-specific lifespan trajectories of spinal cord morphometry. The top panel displays sex-stratified centile curves (5th, 50th, 95th) for Cross-Sectional Area (CSA) across all seven cervical levels (C1–C7). The bottom panel shows the trajectories for all six morphometric features at the representative C3 level. In all plots, raw data points and fitted centiles are shown for males (blue) and females (red). The charts show larger cord size in males and provide sex-specific reference distributions for individual benchmarking.

### Brain-Spinal Cord Relationships

To place spinal cord development and aging within the context of the entire central nervous system, we compared its lifespan trajectory with key brain volumes (**Figure 7A**). The timing of spinal cord growth, maturation, and aging most closely mirrors the lifespan trajectories of total white matter and brainstem volume, with gray matter, thalamic, and total brain volumes peaking earlier in the lifespan. Lifespan-stage correlations (**Figure 7B–C**) are positive and strongest in childhood, decline from rostral to caudal levels, and are most robust for brainstem (followed by white matter). Shape metrics show little to no association with brain volumes. Correlation analyses excluded infancy where brain volumetry did not meet QC and age-stage cells with insufficient sample size.

**Figure 7.**
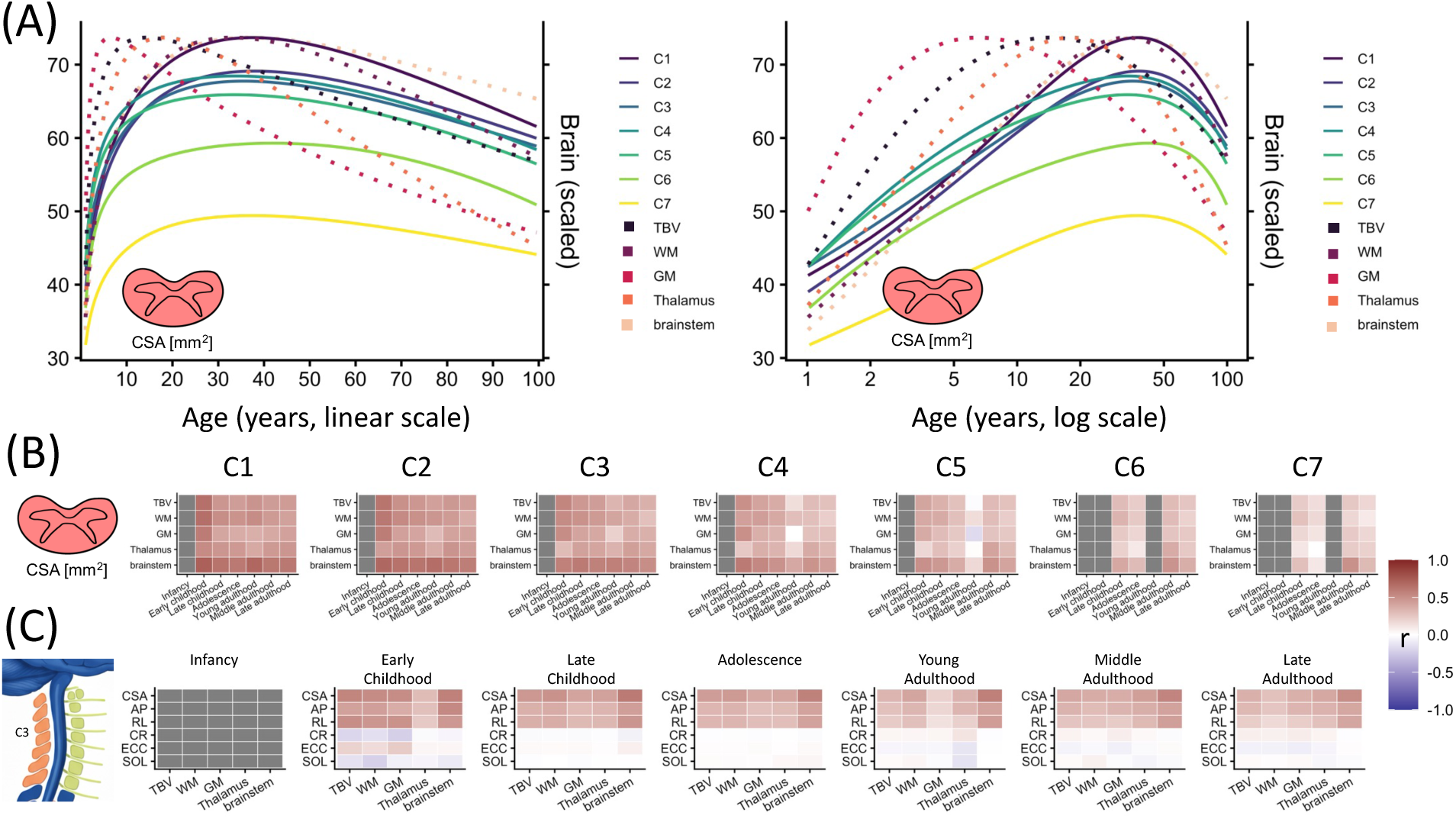
Brain-spinal cord relationships across the lifespan. (A) The median (50th centile) lifespan trajectories for spinal cord Cross-Sectional Area (CSA) at all cervical levels are plotted alongside key brain morphometric volumes. Trajectories are shown on both a linear and a log-scale age axis to accentuate developmental and aging patterns. (B) Pearson correlation coefficients between CSA and brain volumes at different lifespan stages are shown at each cervical level (C1–C7). (C) Correlations between all spinal cord features (at the representative C3 level) and brain volumes are also stratified by lifespan stage. Correlation analyses excluded infancy, as brain volumetric measurements were not consistently reliable in this age range, and any age bin with fewer than 50 subjects. Abbreviations: TBV, total brain volume; WM, total white matter volume; GM, total gray matter volume. We note x-limit is constrained to >0.5 years so that dynamic contrast across the lifespan is visible.

### Aligning new data to the charts

Alignment of new data to the charts is shown in **Figure 8**. Although VUIIS-MS showed only a modest initial offset (**Figure 8A,B**), alignment (**Figure 8C,D**) yielded mean and median control centiles of 50.4 and 48.8, respectively, with controls broadly distributed across the centile range. The same transformation provided individual centiles for participants with MS, whose mean and median centiles were lower (35.4 and 38.3), enabling subsequent analyses of clinical associations and longitudinal change relative to a normative population. A positive-control experiment with an introduced 5% bias confirmed correction of a known dataset offset (**Supplementary Figure 4**), while the larger ADNI example demonstrated alignment and centile extraction in a dataset with a different sample size, population, and acquisition - showing reduced SC CSA in individuals with Alzheimer’s Disease, with 14% of individuals below the 5^th^ centile (**Supplementary Figure 5**).

**Figure 8.**
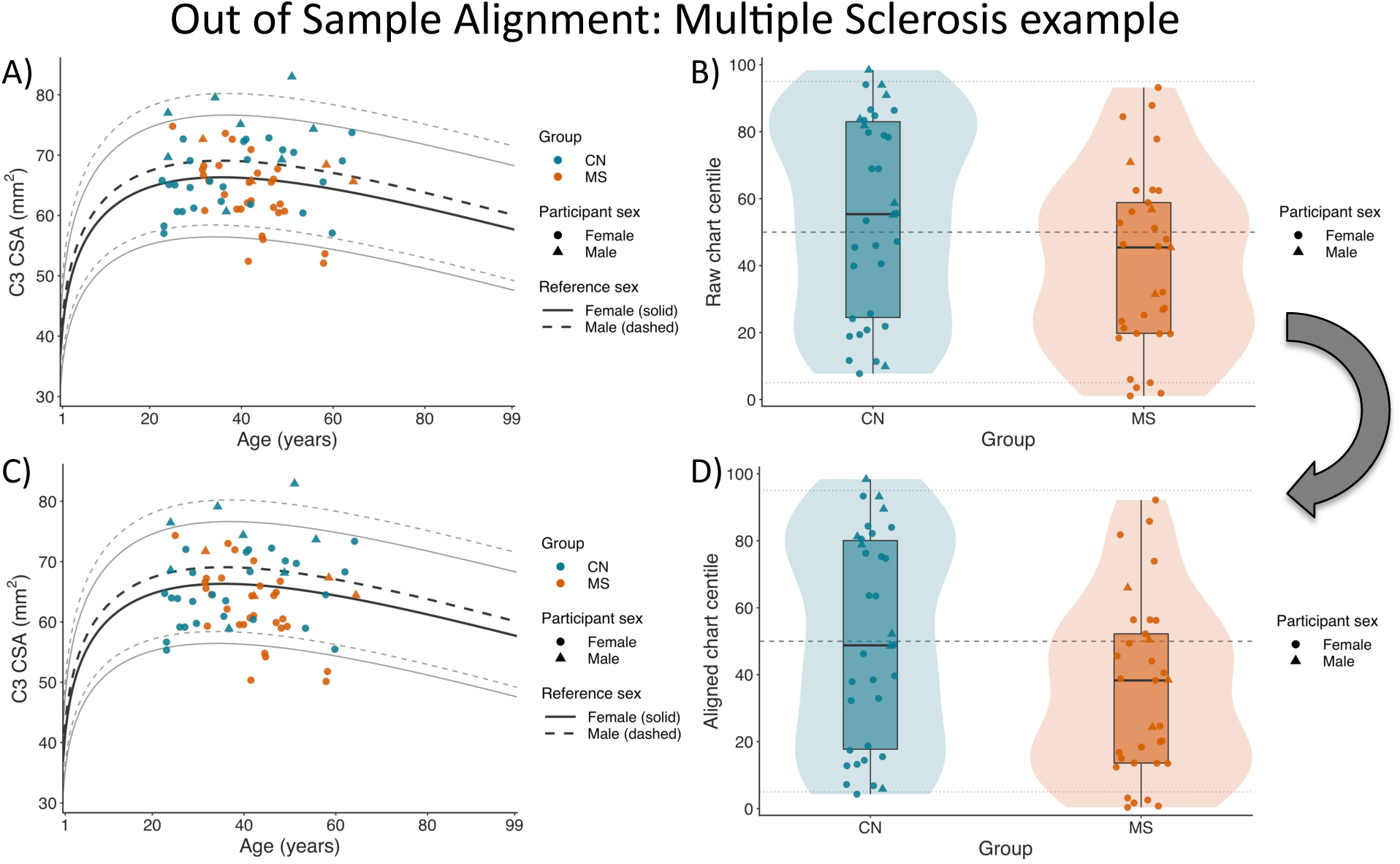
Out-of-sample alignment of a new dataset to the spinal cord charts. The example dataset contained 34 controls and 33 participants with multiple sclerosis (MS) and was not used to build the C3 reference chart used for this analysis. (A) Observed C3 spinal cord cross-sectional area (CSA) plotted against the age- and sex-specific reference chart. (B) Distributions of raw chart centiles in controls and participants with MS. (C) Harmonized C3 CSA after generalized gamma location and scale offsets were estimated from controls and applied unchanged to both groups. (D) Distributions of aligned chart centiles. Reference curves show the 5th, 50th, and 95th centiles; solid curves represent females and dashed curves males. The curved arrow indicates the transformation from raw to aligned measurements.

### Model stability and robustness

Repeated scan selection and leave-one-dataset-out refits produced closely overlapping median centile trajectories (**Supplementary Figures 6–11**). Across features, sexes, and ages at C1– C5, the median CV was 0.05% across repeated scan selections (95th percentile, 0.23%) and 0.12% across leave-one-dataset-out refits (95th percentile, 0.33%); variability was greater at C6–C7 and sparsely sampled ages.

## Discussion

We generated vertebral-level, sex-stratified normative charts for cervical spinal cord morphometry in 24,251 healthy or cognitively normal controls from 33 datasets spanning ages 0-100 years, using contrast-agnostic segmentation [2] and statistical lifespan modeling [42] to harmonize measurements. These charts model the full distribution of CSA, diameters (AP, RL), and shape indices (AP/RL ratio, eccentricity, solidity) across the lifespan, revealing non-linear trajectories - rapid early growth, a mid-adult peak (mid-30s for CSA), and gradual decrease - with clear level-wise differences and stable, low relative dispersion. Together with sex-stratified curves and brain-cord comparisons, the resource provides a population reference that addresses long-standing inter-individual variability [23 and enables age- and sex-adjusted interpretation of individual measurements.

### Spinal Cord Charts as a Tool for Normative Assessment

Normative reference charts for the spinal cord may potentially be a powerful new tool for clinical and research applications. These charts translate absolute spinal cord measurements into age- and sex-adjusted centile scores, directly paralleling pediatric growth and recent brain charts [31, 46]. A centile expresses where an individual lies within the healthy distribution at their age/sex, clarifying whether a small CSA reflects pathology or the lower tail of typical variation. In clinical contexts, a value below a prespecified centile (e.g., 5th) may flag atypical caliber even when absolute thresholds are ambiguous, improving interpretability for disorders where cord atrophy is relevant.

Clinically, this framing helps in scenarios where variability obscures signal: (i) early disease (e.g., suspected inflammatory or neurodegenerative cord involvement) where a patient’s CSA or diameter is well below age-/sex-expected centiles despite modest absolute deviations; (ii) heterogeneous datasets (different scanners/protocols) where centiles provide a common interpretive scale; and (iii) monitoring where a drop across centiles over time suggests accelerated decline beyond age-expected change. In each case, centiles might complement radiological assessment and absolute metrics, adding standardized context to individual measurements.

For research and trials, centile scores offer variance reduction (age/sex accounted for by design) and standardized endpoints to compare groups or track change relative to normative trajectories. Trial planners can use the stage-specific annualized rates of change (acknowledging that these were derived from cross-sectional data) to set realistic progression expectations and power calculations, while analyses can test whether interventions preserve centiles (or slow centile decline) versus control. In routine studies, centiles provide a common scale across sites and protocols, supporting reproducible cross-cohort comparisons.

### Using the spinal cord charts

To facilitate practical application, we favored simplicity and accessibility; analogous to pediatric growth charts, we provide reference tables (CSVs) containing sex-stratified centiles at 0.5-year intervals for every morphometric feature and vertebral level. For a new individual, a feature of interest can be derived and instantly and intuitively placed at the nearest age- and sex-adjusted centile. Critically, valid comparison requires that features be derived using the same processing pipeline and version to avoid methodological bias. While not a standalone diagnostic tool, this “centile scoring” enables the visualization of an individual’s development or aging relative to the population, facilitating the identification of trends or extreme deviations (e.g., <5TH or >95th percentiles).

For dataset-level applications with suitable controls, however, we recommend control-referenced out-of-sample alignment, which accounts for dataset-specific location and scale differences before centile assignment, as in prior brain-charting studies [31, 38]. We demonstrated this approach in the held-out VUIIS-MS dataset, where participants with MS occupied descriptively lower C3 centiles than controls (Figure 8); in a positive-control experiment that corrected an introduced 5% measurement bias (**Supplementary Figure 4**); and in a larger held-out ADNI cohort, where participants with Alzheimer disease occupied lower centiles and were more frequently below the 5th centile (**Supplementary Figure 5**).

This resource builds upon previous normative efforts in the literature, representing a shift in scale and scope. Previous work by Frostell et al. [3] provided essential aggregate reviews of segmental diameters (primarily from postmortem examinations), while Papinutto et al. [22, 23, 47] advanced the field by establishing variability and normalization strategies in targeted adult and pediatric cohorts. Most recently, Valošek et al. [21] introduced a high-resolution database of healthy controls (N=203) standardized to the PAM50 template space. Our work complements these efforts by prioritizing sample size and lifespan coverage. Whereas Valošek et al. maximized spatial precision (slice-wise analysis in template space) in a focused cohort, we leveraged vertebral-level aggregation to maximize statistical power across the entire human lifespan (N=24,251; ages 0–100). This trade-off sacrifices some spatial granularity for the robust modeling of age-related variance. Future work may aim to bridge these approaches, potentially mapping our large-scale lifespan variance into the PAM50 coordinate system to combine high spatial fidelity with robust, data-driven lifespan norms.

Crucially, while these charts define population-level expectations, establishing the functional relevance of individual deviations remains a priority for future validation. Emerging evidence links cervical atrophy directly to clinical outcomes, including respiratory dysfunction in ALS [9], cognitive decline in Alzheimer’s disease [10], and irreversible motor disability in MS [8] - suggesting that ‘centile scoring’ may eventually serve as a predictive biomarker for functional impairment.

### Biological Insights into Lifespan Dynamics of the Spinal Cord

Our lifespan charts provide insights into the biological processes underlying spinal cord development, maturation, and aging. By quantitatively mapping the development trajectory, they revealed rapid growth in early life followed by a protracted maturation extending well into early adulthood [34]. We observe that CSA and cord diameters continue to increase through adolescence, reaching peak values around the mid-30s for CSA. This timeline suggests that the structural maturation of the human spinal cord persists considerably longer than puberty, likely driven by ongoing myelination and axonal growth during young adulthood. Indeed, extensive literature demonstrates that myelination within the central nervous system, in all association, projection, and commissural pathways, continues into the third and even fourth decades of life [48–54]. Our finding of a peak CSA in the mid-30s refines previous estimates from smaller adult cohorts which suggested a peak slightly later, around 45 years [23], potentially reflecting the enhanced temporal resolution afforded by our large-scale dataset.

Following this peak, our charts quantify gradual reductions in cord size with aging; AP diameter declined more rapidly than RL diameter at every cervical level in late adulthood, reversing the greater RL growth seen in early childhood. This imbalance was generally accompanied by a small decrease in AP/RL ratio and increase in eccentricity, while solidity changed little. CSA and diameters are size-based measures commonly interpreted in relation to atrophy, whereas the shape indices describe geometry and should not themselves be interpreted as direct measures of tissue loss. The underlying neurobiology of this late-life cord atrophy likely involves a multifactorial process analogous to brain aging, including progressive demyelination, axonal loss, and neuronal shrinkage or death in spinal gray matter. This reinforces the view of the spinal cord as an integral part of the CNS aging process and suggests its metrics can act as a complement to brain measures. In summary, our charts delineate key biological milestones - the age of peak maturation and the onset and rate of age-related decline - providing a quantitative framework for understanding spinal cord structure across the lifespan.

### An Integrated Central Nervous System: Brain–Cord Relationships and Sex Differences

Our findings reinforce the view of the central nervous system developing and aging as an integrated unit [55, 56], highlighting both coordinated brain-cord dynamics and sex differences. We find strong positive cross-sectional correlations between cervical cord CSA and global brain volumes (total brain, white matter, gray matter, thalamus, brainstem)[22, 25, 57], particularly during developmental periods (**Figure 7B, C**). This brain–cord coupling suggests an intuitive scaling effect: individuals with larger brains tend to have proportionally larger spinal cords, likely reflecting shared genetic and developmental influences on overall nervous system size [25].

This relationship motivates normalization strategies that adjust cord measures for intracranial or brain volume to reduce inter-subject variability [1, 5, 6, 22, 23]. Such approaches may improve sensitivity in some applications, but produce a different reference and require a usable brain image. Further, the lifespan trajectories show spinal cord morphometry changes closely paralleling those of brain white matter and brainstem volume (**Figure 7A**). This indicates that the spinal cord does not age in isolation but rather undergoes senescence in concert with the brain, again suggesting shared underlying biological mechanisms [14]. In pathological states like MS, this link is often more pronounced, with concurrent brain and spinal cord atrophy reflecting widespread neurodegeneration [15, 58, 59]. Understanding this normative brain-cord relationship opens possibilities for developing combined brain-plus-cord centile scores as more holistic biomarkers of CNS health, aging, or disease burden, potentially identifying individuals where one component deviates unexpectedly relative to the other.

We also confirmed sex differences in spinal cord morphometry, necessitating the use of sex-stratified normative charts (**Figure 6**). Males exhibited consistently larger cervical CSA and diameters than females across the lifespan. This sexual dimorphism likely reflects known differences in overall body size, spinal canal dimensions, and potentially hormonal influences on neural development. Practically, this means that comparing an individual’s measurement to sex-specific norms is necessary for accurate assessment and avoids misclassification due to inherent size differences. Our sex-stratified charts provide this calibration, ensuring appropriate benchmarking for both males and females throughout life. The present models generate sex-specific centile curves through fixed sex effects on distribution location and scale; future work should test whether age-dependent changes in location, variability, or skewness differ by sex.

### Limitations and Future Directions

While these first lifespan charts represent a significant advance for spinal cord morphometry, several limitations should be acknowledged. First, our models are derived from cross-sectional data, which approximates lifespan trajectories but cannot capture individual longitudinal changes or potential cohort effects [60]. Although our approach mirrors recent large-scale brain charting initiatives [31], future integration of longitudinal data is needed to fully validate these normative curves. However, our stability analyses provide confidence in the current models: sensitivity analyses support the stability of the modeling framework, demonstrating low uncertainty across alternative scan selections and robustness to the exclusion of individual datasets (Supplementary Figures 6–11) (**Supplementary Figures 6-11**).

Further limitations include the uneven sampling across age groups and lower sample sizes at caudal levels C6 and C7 (**Figure 1**), which reduces model precision at the extremes of the age range and lower cervical spine. For example, C7 had few (N=33) individuals between 20-40 years of age. Moreover, because the cervical cord lies at the periphery of the field of view in standard brain MRI protocols, measurements at these caudal levels are susceptible to gradient non-linearity distortions, a known source of morphometric variability that is particularly pronounced at the edges of the scanner field. This technical confound likely contributes to the larger coefficient of variation observed at C7 compared to rostral levels, rendering maturation and peak estimates less precise in this region. Infancy was sparsely sampled and showed elevated variability for some caudal measures; therefore, both infancy rates of change and variability estimates should be interpreted cautiously. Fitting with a log-age range was tested, with similar results. Future work should include a more dense sampling of early and late-infancy. We advocate caution and consideration when using our charts for C6/C7 levels, and for early infancy (<1 years of age).

Methodologically, our charts are anchored to *vertebral* levels, a practical MRI landmark, rather than functional *neuronal* segments. This is particularly relevant for lifespan modeling because the spine and spinal cord do not grow at the same rate, causing the spatial correspondence between vertebral levels and underlying spinal segments to shift with age. Because of this, observed age-related changes in morphometry at a fixed vertebral level could partially reflect this shifting anatomical frame of reference rather than solely intrinsic tissue changes. While vertebral referencing aligns with common clinical practice, and is practically feasible with noninvasive clinical MRI, future work could map these norms onto segmental anatomy [3, 61] or use developmentally stable landmarks like the pontomedullary junction [1]. Related to modeling, the fractional-polynomial specification is robust but may underfit abrupt developmental transitions or more complex, multimodal lifespan trajectories. Similarly, holding the generalized gamma shape parameter constant across age, sex, and dataset may not capture changes in distributional skewness and may reduce accuracy at the extreme centiles.

Each feature was modeled independently to retain simple, interpretable centiles for individual measurements. Future work could model covariance across measures and vertebral levels to derive lower-dimensional morphometric profiles. Next, our charts are restricted to cervical cord due to the nature of repurposed brain imaging datasets; future work can extend charting to the thoracic and lumbar cord using dedicated spine datasets. The normative values are also inherently tied to the state-of-the-art contrast-agnostic segmentation method employed; application to data processed with different algorithms may require calibration and/or harmonization. Next, averaging across vertebral levels versus slice-wise (e.g. in PAM50 space, as in [21]) may overshadow more subtle changes happening within a level or at disc levels (i.e., stenosis). Moreover, we covary for age and sex, but other factors could explain variance – ethnicity, size, neck length, etc, and it remains to be seen how best to integrate these phenotypes to identify typical versus atypical spinal cord. Finally, while site effects were statistically modeled, residual variance due to scanner hardware or specific protocols might persist [26, 27].

The crucial next step is validating the clinical utility of these charts by applying centile scoring in patient populations (e.g., MS, SCI, DCM) to determine their sensitivity in detecting atypical development or pathological atrophy. Integrating these spinal cord charts with existing brain charts could also enable powerful, composite biomarkers for assessing overall CNS health and disease.

## Conclusion

By using large-scale brain imaging cohorts with a contrast-agnostic segmentation approach, we have generated the first comprehensive normative charts detailing the non-linear dynamics of cervical spinal cord morphometry across the human lifespan. These charts provide reference data on typical growth, maturation, aging, sex differences, and population variability, revealing parallel development with brain structures like white matter and the brainstem. Analogous to established growth charts, these spinal cord charts offer a framework for converting individual measurements into age- and sex-adjusted centile scores, and a standardized tool to benchmark spinal cord health and detect atypical variations indicative of neurodevelopmental or neurodegenerative processes. This work lays the foundation for more precise, individualized assessment of spinal cord integrity in clinical practice and research, with the potential to enhance diagnostic accuracy, improve statistical power in clinical trials, and deepen our understanding of central nervous system development and aging. Future extensions incorporating longitudinal data, thoracic/lumbar regions, and clinical populations will further refine these essential neuroanatomical resources. The lifespan charts, model fits, and out-of-sample alignment code have been made available at (10.5281/zenodo.18306654) to facilitate the use and application of these charts.

## Acknowledgements & Funding

This work was supported in part by the National Institute of Health through NIH awards K01-EB032898 (Schilling) and K01-AG073584 (Archer), grant number 1R01EB017230-01A1 (Landman), K24-AG046373 (Jefferson), and ViSE/VICTR VR3029, UL1-TR000445, and UL1-TR002243. This work was supported by NIA grants R01-AG034962 (Jefferson), R01-AG056534 (Jefferson), R01-AG062826, Alzheimer’s Association IIRG-08-88733 (Jefferson), UL1-TR000445 and UL1-TR002243 (Jefferson), and P20-AG068082 (Jefferson), K01MH090232, R21MH101321, R01MH102272, and the NICHD, R01-HD114489 (Vinci-Booher). This work was supported by the Alzheimer’s Disease Sequencing Project Phenotype Harmonization Consortium (ADSP-PHC) that is funded by NIA (U24 AG074855, U01 AG068057 and R01 AG059716). This work was conducted in part using the resources of the Advanced Computing Center for Research and Education (ACCRE) at Vanderbilt University, Nashville, TN. We appreciate the National Institute of HealthS10 Shared Instrumentation grant 1S10OD020154-01, grant 1S10OD023680-01 (Vanderbilt’s High-Performance Computer Cluster for Biomedical Research), and S10-OD021771 (VUIIS Center for Human Imaging). This work was also supported in part by Intramural Research Program of the National Institute on Aging, NIH. Research reported in this publication was supported by NIGMS of the National Institutes of Health under award number T32GM007347 and T32GM152284.

JV received funding from the European Union’s Horizon Europe research and innovation programme under the Marie Skłodowska-Curie grant agreement No 101107932.

This research was supported in part by the Intramural Research Program of the National Institutes of Health (NIH). The contributions of the NIH author(s) were made as part of their official duties as NIH federal employees, are in compliance with agency policy requirements, and are considered Works of the United States Government. However, the findings and conclusions presented in this paper are those of the author(s) and do not necessarily reflect the views of the NIH or the U.S. Department of Health and Human Services.

BABIES and ABC datasets were supported by the Jacobs Foundation Early Career Research Fellowship (2017-1261-05) (Humphreys); National Science Foundation CAREER Award (2042285) (Humphreys); Brain and Behavior Research Foundation John and Polly Sparks Foundation Investigator Award (29593) (Humphreys); Vanderbilt Institute for Clinical and Translational Research Grant (VR53419) (Humphreys); Vanderbilt Strong Grant; Vanderbilt Kennedy Center Grant (Humphreys); National Institute of Mental Health (R01MH129634) (Humphreys).

This work was supported by the NIH NINDS through award numbers R01 NS108445 and R01 NS110130 (Morgan), and R01 NS134625 and R01 NS112252 (Englot). Data collected from the MORGAN dataset were acquired at Vanderbilt University, with the aim of studying cognitive patterns in patients with epilepsy both before and after clinical treatment.

This work was supported by the NIH Eunice Kennedy Shriver National Institute Of Child Health & Human Development through grant number F31 HD104385 (Nguyen), P50HD103537 (PI: Neul), R37 HD095519 MERIT Award, R01 HD089474, R01 HD109151, R01 HD067254, and R01 HD044073 (Cutting). This work was also supported by the NIH NINDS through grant number R01 NS049096 (Cutting).

This work was supported by the Canada Research Chair in Quantitative Magnetic Resonance Imaging [CRC-2020-00179], the Canadian Institute of Health Research [PJT-190258, PJT-203803], the Canada Foundation for Innovation [32454, 34824], the Fonds de Recherche du Québec - Santé [322736, 324636], the Natural Sciences and Engineering Research Council of Canada [RGPIN-2019-07244], the Canada First Research Excellence Fund (IVADO and TransMedTech), the Courtois NeuroMod project, the Quebec BioImaging Network [5886, 35450], Mila - Tech Transfer Funding Program.

Data and/or research tools used in the preparation of this manuscript were obtained from the National Institute of Mental Health (NIMH) Data Archive (NDA). NDA is a collaborative informatics system created by the National Institutes of Health to provide a national resource to support and accelerate research in mental health. This manuscript reflects the views of the authors and may not reflect the opinions or views of the NIH or of the Submitters submitting original data to NDA.

The Pediatric Imaging, Neurocognition, and Genetics (PING) dataset was collected and released openly to contribute to the assessment of typical brain development in a pediatric sample (RC2DA029475-01), (https://www.sciencedirect.com/science/article/pii/S1053811915003572).

The data used in this study come from the Human Connectome Project, which aims to map the structural connections and circuits of the brain and their relationships to behavior by acquiring high-quality magnetic resonance images. We used diffusion MRI data from the Human Connectome Project Aging (HCPA) study.

The Healthy Brain Network (HBN) is an ongoing initiative focused on building a biobank of data from 10,000 children and adolescents (ages 5-21) in the New York City area (https://www.nature.com/articles/sdata2017181). Data were acquired from the DSI studio website: (https://brain.labsolver.org/hbn.html).

Data collection and sharing for this project was provided by the Cambridge Centre for Ageing and Neuroscience (CamCAN). CamCAN funding was provided by the UK Biotechnology and Biological Sciences Research Council (grant number BB/H008217/1), together with support from the UK Medical Research Council and University of Cambridge, UK. Data used in the preparation of this work were obtained from the CamCAN repository (available at http://www.mrc-cbu.cam.ac.uk/datasets/camcan/). The Cambridge Centre for Ageing and Neuroscience (Cam-CAN) is a large-scale collaborative research project at the University of Cambridge.

The Vanderbilt Memory and Aging Project (VMAP) and the Tennessee Alzheimer’s Project (TAP) data were collected by the Vanderbilt Memory and Alzheimer’s Center (VMAC) and the Vanderbilt Alzheimer’s Disease Research Center’s (VADRC) Investigators at Vanderbilt University Medical Center. VMAP began in 2012 with the goal of investigating vascular health and brain aging. TAP began in 2020 as part of the Exploratory VADRC (P20), which, since 2025, has been established as a Center of Research Excellence (P30).

BLSA is a prospective cohort study with continuous enrollment that began in 1958. Comprehensive data from BLSA are available upon request by a proposal submission through the cohort website (www.blsa.nih.gov). The BLSA is supported by the Intramural Research Program of the National Institute on Aging, NIH. The contributions of the NIH authors are considered Works of the United States Government. The findings and conclusions presented in this paper are those of the authors and do not necessarily reflect the views of the NIH or the U.S. Department of Health and Human Services.

The BIOCARD study is designed to identify biomarkers associated with progression from normal cognitive status to cognitive impairment or dementia, with a particular focus on Alzheimer’s Disease.

Data collection and sharing for this project was funded by the Alzheimer’s Disease Neuroimaging Initiative(ADNI) (National Institutes of Health Grant U01 AG024904) and DOD ADNI (Department of Defense award number W81XWH-12-2-0012). ADNI is funded by the National Institute on Aging, the National Institute of Biomedical Imaging and Bioengineering, and through generous contributions from the following: AbbVie,Alzheimer’s Association; Alzheimer’s Drug Discovery Foundation; Araclon Biotech; BioClinica, Inc.; Biogen; Bristol-Myers Squibb Company; CereSpir, Inc.; Cogstate; Eisai Inc.; Elan Pharmaceuticals, Inc.; Eli Lilly and Company; EuroImmun; F. Hoffmann-La Roche Ltd and its affiliated company Genentech, Inc.; Fujirebio; GEHealthcare; IXICO Ltd.; Janssen Alzheimer Immunotherapy Research & Development, LLC.; Johnson &Johnson Pharmaceutical Research & Development LLC.; Lumosity; Lundbeck; Merck & Co., Inc.; Meso Scale Diagnostics, LLC.; NeuroRx Research; Neurotrack Technologies; Novartis Pharmaceuticals Corporation; Pfizer Inc.; Piramal Imaging; Servier; Takeda Pharmaceutical Company; and Transition Therapeutics. The Canadian Institutes of Health Research is providing funds to support ADNI clinical sites in Canada. Private sector contributions are facilitated by the Foundation for the National Institutes of Health(www.fnih.org). The grantee organization is the Northern California Institute for Research and Education, and the study is coordinated by the Alzheimer’s Therapeutic Research Institute at the University of Southern California. ADNI data are disseminated by the Laboratory for Neuro Imaging at the University of Southern California.

HABS-HD research data reported on this publications was supported by the National Institute on Aging of the National Institutes of Health under Award Numbers R01AG054073, R01AG058533, R01AG070862, P41EB015922 and U19AG078109. The content is solely the responsibility of the authors and does not necessarily represent the official views of the National Institutes of Health.

The NACC database is funded by NIA/NIH Grant U24 AG072122. NACC data are contributed by the NIA-funded ADRCs: P30 AG062429 (PI James Brewer, MD, PhD), P30 AG066468 (PI Oscar Lopez, MD), P30 AG062421 (PI Bradley Hyman, MD, PhD), P30 AG066509 (PI Thomas Grabowski, MD), P30 AG066514 (PI Mary Sano, PhD), P30 AG066530 (PI Helena Chui, MD), P30 AG066507 (PI Marilyn Albert, PhD), P30 AG066444 (PI John Morris, MD), P30 AG066518 (PI Jeffrey Kaye, MD), P30 AG066512 (PI Thomas Wisniewski, MD), P30 AG066462 (PI Scott Small, MD), P30 AG072979 (PI David Wolk, MD), P30 AG072972 (PI Charles DeCarli, MD), P30 AG072976 (PI Andrew Saykin, PsyD), P30 AG072975 (PI David Bennett, MD), P30 AG072978 (PI Ann McKee, MD), P30 AG072977 (PI Robert Vassar, PhD), P30 AG066519 (PI Frank LaFerla, PhD), P30 AG062677 (PI Ronald Petersen, MD, PhD), P30 AG079280 (PI Eric Reiman, MD), P30 AG062422 (PI Gil Rabinovici, MD), P30 AG066511 (PI Allan Levey, MD, PhD), P30 AG072946 (PI Linda Van Eldik, PhD), P30 AG062715 (PI Sanjay Asthana, MD, FRCP), P30 AG072973 (PI Russell Swerdlow, MD), P30 AG066506 (PI Todd Golde, MD, PhD), P30 AG066508 (PI Stephen Strittmatter, MD, PhD), P30 AG066515 (PI Victor Henderson, MD, MS), P30 AG072947 (PI Suzanne Craft, PhD), P30 AG072931 (PI Henry Paulson, MD, PhD), P30 AG066546 (PI Sudha Seshadri, MD), P20 AG068024 (PI Erik Roberson, MD, PhD), P20 AG068053 (PI Justin Miller, PhD), P20 AG068077 (PI Gary Rosenberg, MD), P20 AG068082 (PI Angela Jefferson, PhD), P30 AG072958 (PI Heather Whitson, MD), P30 AG072959 (PI James Leverenz, MD).

The NACC database is funded by NIA/NIH Grant U24 AG072122. SCAN is a multi-institutional project that was funded as a U24 grant (AG067418) by the National Institute on Aging in May 2020. Data collected by SCAN and shared by NACC are contributed by the NIA-funded ADRCs as follows:

Arizona Alzheimer’s Center - P30 AG072980 (PI: Eric Reiman, MD); R01 AG069453 (PI: Eric Reiman (contact), MD); P30 AG019610 (PI: Eric Reiman, MD); and the State of Arizona which provided additional funding supporting our center; Boston University - P30 AG013846 (PI Neil Kowall MD); Cleveland ADRC - P30 AG062428 (James Leverenz, MD); Cleveland Clinic, Las Vegas – P20AG068053; Columbia - P50 AG008702 (PI Scott Small MD); Duke/UNC ADRC – P30 AG072958; Emory University - P30AG066511 (PI Levey Allan, MD, PhD); Indiana University - R01 AG19771 (PI Andrew Saykin, PsyD); P30 AG10133 (PI Andrew Saykin, PsyD); P30 AG072976 (PI Andrew Saykin, PsyD); R01 AG061788 (PI Shannon Risacher, PhD); R01 AG053993 (PI Yu-Chien Wu, MD, PhD); U01 AG057195 (PI Liana Apostolova, MD); U19 AG063911 (PI Bradley Boeve, MD); and the Indiana University Department of Radiology and Imaging Sciences; Johns Hopkins - P30 AG066507 (PI Marilyn Albert, Phd.); Mayo Clinic - P50 AG016574 (PI Ronald Petersen MD PhD); Mount Sinai - P30 AG066514 (PI Mary Sano, PhD); R01 AG054110 (PI Trey Hedden, PhD); R01 AG053509 (PI Trey Hedden, PhD); New York University - P30AG066512-01S2 (PI Thomas Wisniewski, MD); R01AG056031 (PI Ricardo Osorio, MD); R01AG056531 (PIs Ricardo Osorio, MD; Girardin Jean-Louis, PhD); Northwestern University - P30 AG013854 (PI Robert Vassar PhD); R01 AG045571 (PI Emily Rogalski, PhD); R56 AG045571, (PI Emily Rogalski, PhD); R01 AG067781, (PI Emily Rogalski, PhD); U19 AG073153, (PI Emily Rogalski, PhD); R01 DC008552, (M.-Marsel Mesulam, MD); R01 AG077444, (PIs M.-Marsel Mesulam, MD, Emily Rogalski, PhD); R01 NS075075 (PI Emily Rogalski, PhD); R01 AG056258 (PI Emily Rogalski, PhD); Oregon Health and Science University - P30 AG008017 (PI Jeffrey Kaye MD); R56 AG074321 (PI Jeffrey Kaye, MD); Rush University - P30 AG010161 (PI David Bennett MD); Stanford – P30AG066515; P50 AG047366 (PI Victor Henderson MD MS); University of Alabama, Birmingham – P20; University of California, Davis - P30 AG10129 (PI Charles DeCarli, MD); P30 AG072972 (PI Charles DeCarli, MD); University of California, Irvine - P50 AG016573 (PI Frank LaFerla PhD); University of California, San Diego - P30AG062429 (PI James Brewer, MD, PhD); University of California, San Francisco - P30 AG062422 (Rabinovici, Gil D., MD); University of Kansas - P30 AG035982 (Russell Swerdlow, MD); University of Kentucky - P30 AG028283-15S1 (PIs Linda Van Eldik, PhD and Brian Gold, PhD); University of Michigan ADRC - P30AG053760 (PI Henry Paulson, MD, PhD) P30AG072931 (PI Henry Paulson, MD, PhD) Cure Alzheimer’s Fund 200775 - (PI Henry Paulson, MD, PhD) U19 NS120384 (PI Charles DeCarli, MD, University of Michigan Site PI Henry Paulson, MD, PhD) R01 AG068338 (MPI Bruno Giordani, PhD, Carol Persad, PhD, Yi Murphey, PhD) S10OD026738-01 (PI Douglas Noll, PhD) R01 AG058724 (PI Benjamin Hampstead, PhD) R35 AG072262 (PI Benjamin Hampstead, PhD) W81XWH2110743 (PI Benjamin Hampstead, PhD) R01 AG073235 (PI Nancy Chiaravalloti, University of Michigan Site PI Benjamin Hampstead, PhD) 1I01RX001534 (PI Benjamin Hampstead, PhD) IRX001381 (PI Benjamin Hampstead, PhD); University of New Mexico - P20 AG068077 (Gary Rosenberg, MD); University of Pennsylvania - State of PA project 2019NF4100087335 (PI David Wolk, MD); Rooney Family Research Fund (PI David Wolk, MD); R01 AG055005 (PI David Wolk, MD); University of Pittsburgh - P50 AG005133 (PI Oscar Lopez MD); University of Southern California - P50 AG005142 (PI Helena Chui MD); University of Washington - P50 AG005136 (PI Thomas Grabowski MD); University of Wisconsin - P50 AG033514 (PI Sanjay Asthana MD FRCP); Vanderbilt University – P20 AG068082; Wake Forest - P30AG072947 (PI Suzanne Craft, PhD); Washington University, St. Louis - P01 AG03991 (PI John Morris MD); P01 AG026276 (PI John Morris MD); P20 MH071616 (PI Dan Marcus); P30 AG066444 (PI John Morris MD); P30 NS098577 (PI Dan Marcus); R01 AG021910 (PI Randy Buckner); R01 AG043434 (PI Catherine Roe); R01 EB009352 (PI Dan Marcus); UL1 TR000448 (PI Brad Evanoff); U24 RR021382 (PI Bruce Rosen); Avid Radiopharmaceuticals / Eli Lilly; Yale - P50 AG047270 (PI Stephen Strittmatter MD PhD); R01AG052560 (MPI: Christopher van Dyck, MD; Richard Carson, PhD); R01AG062276 (PI: Christopher van Dyck, MD); 1Florida - P30AG066506- 03 (PI Glenn Smith, PhD); P50 AG047266 (PI Todd Golde MD PhD) Data contributed from MAP/ROS/MARS was supported by NIA R01AG017917, P30AG10161, P30AG072975, R01AG022018, R01AG056405, UH2NS100599, UH3NS100599, R01AG064233, R01AG15819 and R01AG067482, and the Illinois Department of Public Health (Alzheimer’s Disease Research Fund). Data can be accessed at www.radc.rush.edu. More information about participant demographics and study information can be found here: https://www.rushu.rush.edu/research-rush-university/departmental-research/rush-alzheimers-disease-center/rush-alzheimers-disease-center-research/epidemiologic-research.

The data contributed from the Wisconsin Registry for Alzheimer’s Prevention was supportedby NIA AG021155, AG0271761, AG037639, and AG054047.

Data collection and sharing for this project was provided by the Centre for Attention, Learning and Memory (CALM). CALM funding was provided by the UK Medical Research Council and University of Cambridge, UK. Data used in the preparation of this work were obtained from CALM resource – https://calm.mrc-cbu.cam.ac.uk/. The study protocol is reported in Holmes et al. (2019).

Data used in the preparation of this work were obtained from the International Consortium for Brain Mapping (ICBM) database (www.loni.usc.edu/ICBM). The ICBM project (Principal Investigator John Mazziotta, M.D., University of California, Los Angeles) is supported by the National Institute of Biomedical Imaging and BioEngineering. ICBM is the result of efforts of co-investigators from UCLA, Montreal Neurologic Institute, University of Texas at San Antonio, and the Institute of Medicine, Juelich/Heinrich Heine University - Germany. Data collection and sharing for this project was provided by the International Consortium for Brain Mapping (ICBM; Principal Investigator: John Mazziotta, MD, PhD). ICBM funding was provided by the National Institute of Biomedical Imaging and BioEngineering. ICBM data are disseminated by the Laboratory of Neuro Imaging at the University of Southern California.

The UCLA Consortium for Neuropsychiatric Phenomics LA5c Study (UCLA) is focused on understanding the dimensional structure of memory and cognitive control (response inhibition) functions in both healthy individuals and individuals with neuropsychiatric disorders including schizophrenia, bipolar disorder, and attention deficit/hyperactivity disorder. Neuroimaging data were downloaded from Openneuro here: https://openneuro.org/datasets/ds000030/versions/1.0.0. We use version 1.0.0.

The Dallas Lifespan Brain Study (DLBS) is a longitudinal multi-modal neuroimaging study of the aging mind, which was initiated in 2008 (referred to as Wave 1). Participants returned for two additional waves of data collection with an approximate interval of 4-5 years between waves. The DLBS protocol encompasses various imaging modalities, including structural MRI, diffusion MRI, and functional MRI, as well as comprehensive cognitive and psychosocial assessments. DLBS data can be downloaded from Openneuro here: https://openneuro.org/datasets/ds004856. Specifically, we use version 1.2.0.

The Multisite, Multiscanner, and Multisubject Acquisitions for Studying Variability in Diffusion Weighted Magnetic Resonance Imaging (MASiVar) dataset consists of 319 diffusion scans acquired at 3T from b = 1000 to 3000 s/mm2 across 14 healthy adults, 83 healthy children (5 to 8 years), three sites, and four scanners curated to promote investigation of diffusion MRI variability. In particular, we used only the data coming from healthy children (Cohort IV) for version 2.0.2 of the dataset. Data are available to download from Openneuro here: https://openneuro.org/datasets/ds003416/versions/2.0.2.

The Amsterdam Open MRI Collection (AOMIC) is a collection of three datasets with multimodal (3T) MRI data including structural (T1-weighted), diffusion-weighted, and (resting-state and task-based) functional BOLD MRI data, as well as detailed demographics and psychometric variables from a large set of healthy participants. All raw data is publicly available from the Openneuro data sharing platform: ID1000: https://openneuro.org/datasets/ds003097, PIOP1: https://openneuro.org/datasets/ds002785, PIOP2: https://openneuro.org/datasets/ds002790. We use version 1.2.1 for ID1000 and 2.0.0 for PIOP1 and PIOP2.

The Boston Adolescent Neuroimaging of Depression and Anxiety (BANDA) is a study of 215 adolescents ages 14-17, 152 of whom had a current diagnosis of a DSM-5 (APA, 2013) anxious and/or depressive disorder. The BANDA study collected a rich dataset of brain, clinical, and cognitive/neuropsychological measures from these adolescent subjects. The dataset is available to download upon request on the NDA.

We thank Knight ADRC for providing neuroimaging data to us (WASHU dataset). The data contributed through WASHU (Knight ADRC) was supported by grant numbers P30 AG066444, P01 AG03991, and P01 AG026276. For WASHU, Clinical Dementia Ratings (CDRs) are obtained from assessments by experienced clinicians trained in the use of the CDR.

The NACC database is funded by NIA/NIH Grant U24 AG072122. SCAN is a multi-institutional project that was funded as a U24 grant (AG067418) by the National Institute on Aging in May 2020. Data collected by SCAN and shared by NACC are contributed by the NIA-funded ADRCs as follows:

Data used in the preparation of this article were obtained from the Adolescent Brain Cognitive Development (ABCD) Study (https://abcdstudy.org), held in the NIMH Data Archive (NDA). This is a multisite, longitudinal study designed to recruit more than 10,000 children age 9-10 and follow them over 10 years into early adulthood. The ABCD Study® is supported by the National Institutes of Health and additional federal partners under award numbers U01DA041048, U01DA050989, U01DA051016, U01DA041022, U01DA051018, U01DA051037, U01DA050987, U01DA041174, U01DA041106, U01DA041117, U01DA041028, U01DA041134, U01DA050988, U01DA051039, U01DA041156, U01DA041025, U01DA041120, U01DA051038, U01DA041148, U01DA041093, U01DA041089, U24DA041123, U24DA041147. A full list of supporters is available at https://abcdstudy.org/federal-partners.html. A listing of participating sites and a complete listing of the study investigators can be found at https://abcdstudy.org/consortium_members/. ABCD consortium investigators designed and implemented the study and/or provided data but did not necessarily participate in the analysis or writing of this report. This manuscript reflects the views of the authors and may not reflect the opinions or views of the NIH or ABCD consortium investigators.

The Bipolar & Schizophrenia Consortium for Parsing Intermediate Phenotypes (BSNIP1; 10.15154/tnzs-a323) dataset (R01MH078113-01, R01MH077852-01, R01MH077851-01, R01MH077945-01, R01MH077862-01) and its renewal (BSNIP2; 10.15154/8v3w-et72) (R01MH103368-01, R01MH103366-01) were collected with the aim to improve diagnosis and clinical management of psychosis by defining biologically based biotypes that better predict symptoms, course, and treatment response across major psychoses.

## Data Availability Statement

The lifespan charts, model fits, and out-of-sample alignment code have been made available at (10.5281/zenodo.18306654) to facilitate the use and application of these charts.

Data used in the preparation of this article were obtained from the Alzheimer’s Disease Neuroimaging Initiative (ADNI) database (adni.loni.usc.edu). The ADNI was launched in 2003 as a public-private partnership, led by Principal Investigator Michael W. Weiner, MD. The primary goal of ADNI has been to test whether serial magnetic resonance imaging (MRI), positron emission tomography (PET), other biological markers, and clinical and neuropsychological assessment can be combined to measure the progression of mild cognitive impairment (MCI) and early Alzheimer’s disease (AD). Data from the Alzheimer’s Disease Neuroimaging Initiative (ADNI) are available upon request from https://adni.loni.usc.edu/. Data from the AOMIC-PIOP1 dataset are freely available for download on OpenNeuro: https://openneuro.org/datasets/ds002785/versions/2.0.0. Data from the AOMIC-PIOP2 dataset are freely available for download on OpenNeuro: https://openneuro.org/datasets/ds002790/versions/2.0.0. Data from the AOMIC-ID1000 dataset are freely available for download on OpenNeuro: https://openneuro.org/datasets/ds003097/versions/1.2.1. Data from the Boston Adolescent Neuroimaging of Depression and Anxiety (BANDA) dataset are available upon request from https://www.humanconnectome.org/study/connectomes-related-anxiety-depression. Data from BIOCARD are available upon request after filling out a data use application: https://www.gaaindata.org/partner/BIOCARD. Data from the Baltimore Longitudinal Study of Aging (BLSA) are available upon request from https://www.blsa.nih.gov/. Data from the Centre for Attention Learning and Memory (CALM) dataset are available upon request from https://calm.mrc-cbu.cam.ac.uk/researchers/. Data from the Cambridge Center for Ageing Neuroscience (CAMCAN) dataset are available upon request from https://camcan-archive.mrc-cbu.cam.ac.uk/dataaccess/. Data from the Dallas Lifespan Brain Study (DLBS) dataset are freely available for download on OpenNeuro: https://openneuro.org/datasets/ds004856/versions/1.2.0. Data from the Health & Aging Brain Study - Health Disparities (HABS-HD) dataset are available upon request from https://apps.unthsc.edu/itr/reports. Imaging data and basic demographic information for the Healthy Brain Network (HBN) dataset are freely available to download from https://fcon_1000.projects.nitrc.org/indi/cmi_healthy_brain_network/. Phenotypic data are available upon request by filling out a data use agreement: https://fcon_1000.projects.nitrc.org/indi/cmi_healthy_brain_network/Phenotypic.html. Data from the Human Connectome Project – Aging (HCPA) dataset are available upon request from https://www.humanconnectome.org/study/hcp-lifespan-aging. Data from the International Consortium for Brain Mapping (ICBM) dataset are available upon request from www.loni.usc.edu/ICBM. Data from the Memory and Aging Project (MAP), Religious Orders Study (ROS), and the Minority Aging Research Study (MARS) datasets are available upon request from https://www.radc.rush.edu/. Data from the Multisite, Multiscanner, and Multisubject Acquisitions for Studying Variability in Diffusion Weighted Magnetic Resonance Imaging (MASiVar) dataset are freely available for download on OpenNeuro: https://openneuro.org/datasets/ds003416/versions/2.0.2. Data from the National Alzheimer’s Coordinating Center (NACC) and the Standardized Centralized Alzheimer’s & Related Dementias Neuroimaging (SCAN) are available upon request from https://naccdata.org/requesting-data/data-request-process. Data from the Pediatric Imaging, Neurocognition, and Genetics dataset (PING) are available for download from the National Institutes of Mental Health data archive upon request from https://nda.nih.gov/. Data from the UCLA Consortium for Neuropsychiatric Phenomics LA5c Study (UCLA) dataset are freely available for download on OpenNeuro: https://openneuro.org/datasets/ds000030/versions/1.0.0. Data from the Vanderbilt Memory and Aging Project (VMAP_JEFFERSON, VMAP_2.0) are available upon request from https://vmacdata.org/vmap/data-requests. Data from the Vanderbilt Alzheimer’s Disease Research Centre’s Tennessee Alzheimer’s Project (TAP) are available from https://vmacdata.org/vadrc-tap. Data from the Wisconsin Registry for Alzheimer’s Prevention (WRAP) are available upon request from https://wrap.wisc.edu/data-requests-2/. Data from the Adolescent Brain Cognitive Development (ABCD) Study are available for download upon request from (https://abcdstudy.org/scientists/data-sharing/). <u>Data from</u> <u>the Bipolar & Schizophrenia Consortium for Parsing Intermediate Phenotypes dataset (BSNIP1) and its renewal (BSNIP2)</u> are available for download from the National Institutes of Mental Health data archive upon request from https://nda.nih.gov/. Vanderbilt University data (MORGAN, BABIES-ABC, CUTTING, VUMC-ASD) subject to third party restrictions. Please contact corresponding author for data requests.

