## Supplementary Figures and Tables for "Charting Cervical Spinal Cord Morphometry Across the Lifespan"

***Supplementary Table 1.*** *Datasets, sample size, and median age.*

| Dataset | Subjects (sessions) | Healthy Control subjects (sessions) | Final chart subjects | Min age | Median age | Max age |
|---|---|---|---|---|---|---|
| dHCPinfant | 271 (272) | 271 (272) | 0 | -0.08 | 0.02 | 0.09 |
| Humphreys | 113 (137) | 113 (137) | 19 | 0.07 | 0.10 | 1.05 |
| UCSDACE | 327 (466) | 108 (158) | 106 | 1.00 | 2.25 | 10.42 |
| MASiVar | 96 (144) | 96 (144) | 96 | 5.40 | 6.80 | 47.00 |
| CALM | 341 (341) | 203 (203) | 196 | 5.50 | 9.25 | 17.92 |
| HBN | 3,619 (3,619) | 1,434 (1,434) | 1,007 | 5.02 | 10.10 | 21.82 |
| ABCD | 11,823 (21,018) | 9,845 (15,597) | 9,231 | 8.97 | 10.73 | 15.62 |
| PING | 765 (765) | 765 (765) | 699 | 2.00 | 12.12 | 21.67 |
| BANDA | 207 (207) | 63 (63) | 62 | 14.00 | 15.17 | 16.83 |
| AOMIC | 842 (842) | 842 (842) | 573 | 18.75 | 22.75 | 26.25 |
| UCLA_LA5c | 260 (260) | 121 (121) | 117 | 21.00 | 28.00 | 50.00 |
| BSNIP2 | 740 (740) | 234 (234) | 224 | 18.00 | 32.00 | 61.00 |
| TempleSocial | 107 (107) | 107 (107) | 88 | 21.00 | 36.50 | 80.00 |
| VUIIS | 47 (72) | 25 (38) | 24 | 23.98 | 40.70 | 64.10 |
| BSNIP1 | 1,074 (1,074) | 368 (368) | 237 | 15.00 | 41.00 | 65.00 |
| ICBM | 173 (173) | 173 (173) | 165 | 19.00 | 44.00 | 80.00 |
| CAMCAN | 633 (633) | 633 (633) | 621 | 18.50 | 55.50 | 88.92 |
| HCPA | 725 (725) | 725 (725) | 674 | 36.00 | 58.25 | 100.00 |
| WRAP | 373 (473) | 181 (251) | 44 | 46.95 | 59.23 | 67.46 |
| DLBS | 464 (957) | 464 (957) | 463 | 21.00 | 64.00 | 97.00 |
| VMAP_2.0 | 327 (355) | 270 (281) | 270 | 50.00 | 64.00 | 90.00 |
| HABSHD | 4,253 (6,488) | 2,662 (4,113) | 2,581 | 50.00 | 65.00 | 90.00 |
| OASIS4 | 660 (673) | 47 (47) | 47 | 47.08 | 67.65 | 85.14 |
| Indiana | 615 (1,040) | 287 (410) | 282 | 18.05 | 68.20 | 93.03 |
| VMAP_TAP | 170 (199) | 52 (63) | 52 | 55.00 | 69.00 | 86.00 |
| BLSA | 1,130 (3,370) | 1,056 (2,789) | 1,032 | 22.40 | 69.60 | 97.70 |
| OASIS3 | 991 (1,685) | 720 (1,292) | 706 | 42.69 | 70.06 | 95.65 |
| NACC | 4,551 (6,070) | 2,305 (2,866) | 1,968 | 22.80 | 70.80 | 95.70 |
| BIOCARD | 212 (504) | 151 (343) | 150 | 34.00 | 71.00 | 91.00 |
| ADNI | 3,117 (12,006) | 1,248 (4,208) | 1,216 | 50.50 | 72.31 | 93.80 |
| SCAN | 2,897 (3,266) | 209 (230) | 202 | 48.22 | 73.08 | 93.93 |
| WASHU | 946 (1,979) | 190 (205) | 183 | 61.64 | 73.50 | 92.10 |
| ROSMAPMARS | 850 (2,053) | 729 (1,715) | 651 | 62.85 | 81.13 | 101.91 |
| ABVIB | 66 (74) | 0 (0) | 0 | — | — | — |
| CIBS_BRAIN2 | 34 (68) | 0 (0) | 0 | — | — | — |

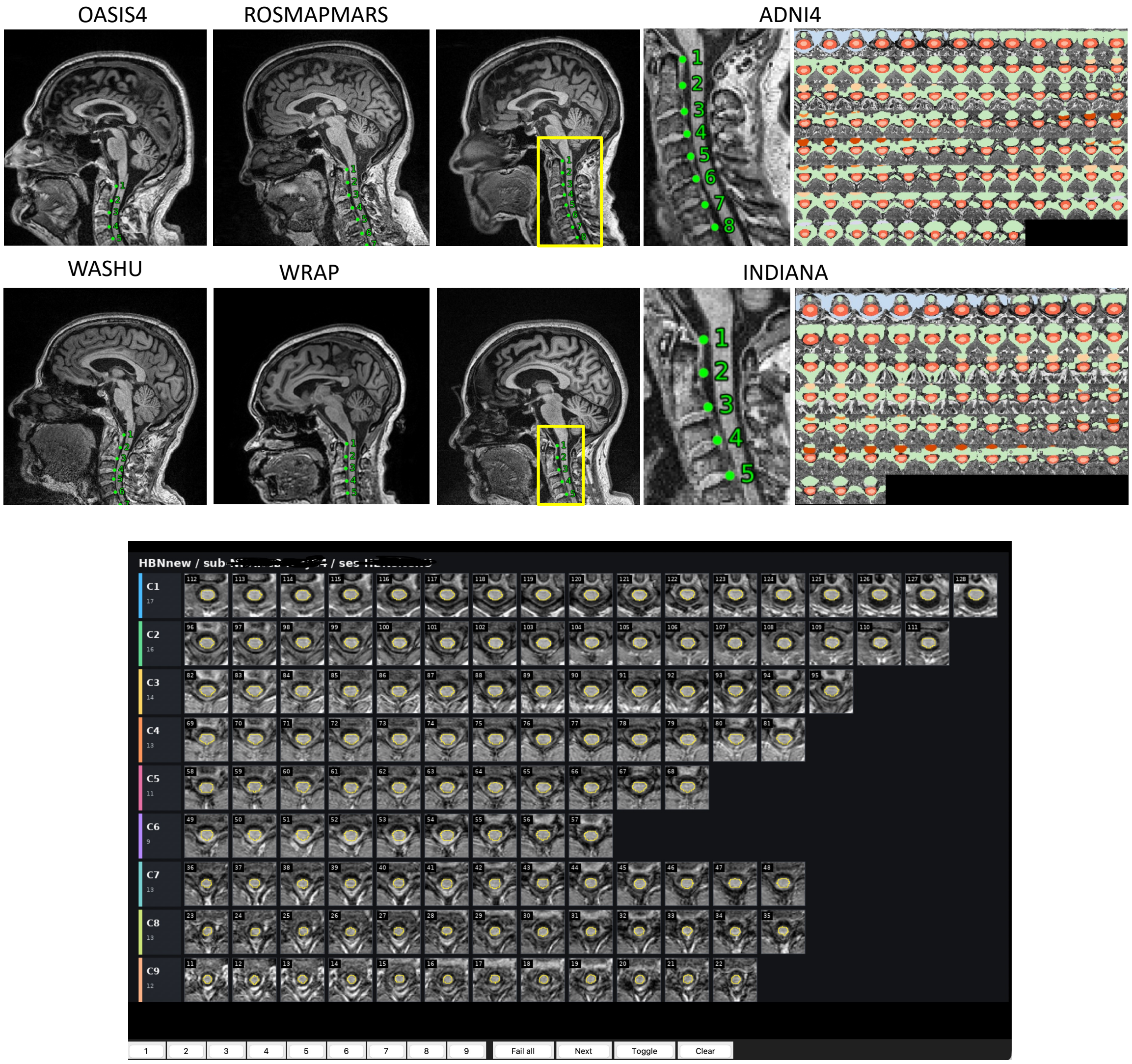


***Supplementary Figure 1.*** *Spinal cord segmentation and vertebral labelling. The spinal cord toolbox contrast-agnostic segmentation is robust to dataset, vendor, and acquisition conditions, with subjects from six cohorts shown (top). Every session was visualized twice (once with vertebral levels, and once with segmentation outline) for quality assurance.*

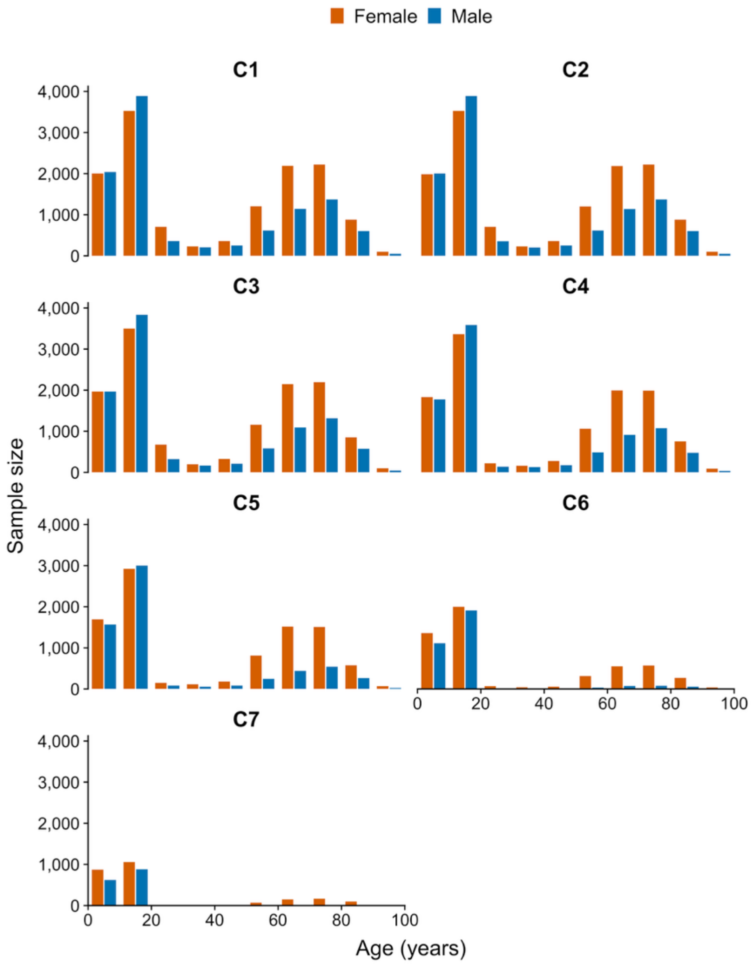


***Supplementary Figure 2***. *Age and sex distributions across cervical levels. Histograms show QA-passing healthy/control participants with a recorded age of 0-100 years in 10-year bins, separated by sex. Participants without recorded age/sex or with age outside 0-100 years are not included. Sample sizes decrease from C1 to C7, with fewer subjects during young adulthood (20-40) and early infancy (<1).*

***Supplementary Table 2**. T1w image cohorts, vendors, sequences.*

| Dataset | Cohort / sites | Vendor | Field strength (T) | T1w sequence | Voxel spacing (mm) | TR (ms) | TE (ms) | TI (ms) | Flip angle (°) |
|---|---|---|---|---|---|---|---|---|---|
| ABCD | 21 sites | GE; Philips; Siemens | 3 | MPRAGE; IR-SPGR; TFE | 1 × 1 × 1 | GE/Siemens 2500†; Philips 6.31† | GE 2.0; Philips 2.9; Siemens 2.88 | 1060 | 8 |
| ADNI | ADNI1 n=10; ADNI-GO n=154; ADNI2 n=323; ADNI3 n=535; ADNI4 | GE; Philips; Siemens | 1.5; 3 | MPRAGE; IR-FSPGR; TFE | ~1 × 1 × 1 to 1 × 1 × 1.2 | 6–9† or 2300–3000† | 2.8–4.0 | 400–1000 | 8–9 |
| AOMIC | ID1000 n=478; PIOP1 n=53; PIOP2 n=42 | Philips | 3 | MPRAGE | 1 × 1 × 1 | ID1000 8.1†; PIOP1/2 8.5† | ID1000 3.7; PIOP1/2 3.9 | NR | 8 |
| BANDA | Single site | Siemens | 3 | vNav MPRAGE | 0.8 × 0.8 × 0.8 | 2400 | 2.18 | 1040 | 8 |
| BIOCARD | Johns Hopkins era | Philips | 3 | MPRAGE | 1 × 1 × 1.2 | 6.75–6.8† | 3.1–3.16 | 843–844 | 8 |
| BLSA | Modern n=1005; legacy n=27 | Legacy: GE/Philips; | 1.5; 3 | SPGR; MPRAGE | Legacy 0.94 × 0.94 × 1.5; modern 1 × 1 × 1.2 | Legacy 35; modern 6.5–6.8† | Legacy 5; modern 3.2 | Legacy NA; modern NR | Legacy 45; modern 8 |
| BSNIP1 | 6 sites | GE; Philips; | 3 | MPRAGE; spoiled GRE | ~1 × 1 × 1.2 | 6.6–7.3† or 2300† | 2.8–3.14 | 650–900 where | 8–9 |
| BSNIP2 | Multisite | GE; Philips; | 3 | MPRAGE; spoiled GRE | ~1 × 1 × 1.2 | 6.72–7.38† or 2300† | 2.83–3.15 | 400–900 where | 9–11 where |
| CALM | Single Cambridge site | Siemens | 3 | MPRAGE | 1 × 1 × 1 | 2250 | 3.02 | 900 | 9 |
| CAMCAN | CC700 Phase 2, Arm 1 | Siemens | 3 | MPRAGE | 1 × 1 × 1 | 2250 | 2.99 | 900 | 9 |
| DLBS | Wave 1 n=291; wave 2 n=106; wave 3 n=66 | Philips | 3 | MPRAGE | 1 × 1 × 1 | 8.08–8.37† | 3.69–3.89 | 1100 | 12 |
| HABS-HD | Baseline n=2138; month 24 n=402; month 48 | Siemens | 3 | MPRAGE | Skyra 1.1 × 1.1 × 1.2; Vida 1 × 1 × 1 | 2300 | Skyra 2.93; Vida 2.98 | Vida 900; Skyra NR | Vida 9; Skyra NR |
| HBN | CBIC n=204; CUNY n=76; Rutgers n=460; Staten Island n=267 | Siemens | 1.5; 3 | MEMPRAGE; MPRAGE | 0.8 × 0.8 × 0.8 or 1 × 1 × 1 | 2500 or 2730 | MEMPRAGE 1.64/3.50/5.36/7.22; MPRAGE 2.88–3.15 | 1000–1070 | 7–8 |
| HCP-Aging | 4 sites | Siemens | 3 | vNav multi-echo MPRAGE | 0.8 × 0.8 × 0.8 | 2500 | 1.8/3.6/5.4/7.2 | 1000 | 8 |
| Humphreys | 6-month n=16; 12-month | Philips | 3 | T1-weighted gradient echo | 0.9375 × 0.9375 × 1 | 8.93–10† | 4.604 | NR | 8 |
| ICBM | MNI n=31; UCLA n=134 | MNI: Siemens; UCLA: GE | 1.5; 3 | SPGR | MNI 1 × 1 × 1; UCLA 1.2 × 1 × 1 | MNI 22; UCLA NR | MNI 9.2; UCLA NR | NA | MNI 30; UCLA NR |
| Indiana | IMAS / Indiana ADRC | Siemens | 3 | MPRAGE | 0.8 × 0.8 × 0.8; 1 × 1 × 1; 1.1 × 1.1 × 1.2 | 2300 | 2.91–2.98 | 900 | 9 |
| VUIIS | Vanderbilt MS cohort | Philips | 3 | T1-weighted gradient echo | 1 × 1 × 1 | 6.31† | 2.90 | NR | 8 |
| MASiVar | Cohort II n=5; cohort III n=8; cohort IV n=83 | Philips; GE; Siemens | 3 | NR | 1 × 1 × 1 | NR | NR | NR | NR |
| NACC | Multiple ADRCs | GE; Philips; Siemens | 1.5; 3 | MPRAGE; FSPGR; BRAVO; TFE; spin echo; | Mixed; commonly ~1 mm | 6.03–450† | 1.8–9.0 | 400–1100 where reported | 7–90 |
| OASIS-3 | Longitudinal; multiple scanner eras | Siemens | 1.5; 3 | MPRAGE | 1 × 1 × 1; ~1.05 × 1.05 × 1.2 | 1900; 2300; 2400 | 3.93; 2.95; 3.16 | NR; 900; 1000 | 15; 9; 8 |
| OASIS-4 | Clinical cohort; multiple protocols | Siemens | 3 | MPRAGE | ~1 × 1 × 1 to 1.055 × 1.055 × 1.1 | 1470–2400 | 2.93–3.47 | 800–1000 | 8–15 |
| PING | Multisite pediatric cohort | GE; Philips; Siemens | 3 | IR-SPGR; TFE; MPRAGE | ~1 × 1 × 1.2 | GE 8.1†; Philips 6.8†; Siemens 2170† | GE 3.5; Philips 3.1; Siemens 2.78 or 4.33 | GE 640; Philips NR; Siemens 1100 | GE/Philips 8; Siemens 7 |
| ROS/MAP/MARS | MAP n=429; MARS n=158; ROS n=64 | Philips; Siemens | 3 | MPRAGE; TFE | 1 × 1 × 1 | Philips 8†; Siemens 2300–2530† | Philips 3.7; Siemens 1.69–2.98 | Philips 955; Siemens | Philips 8; Siemens 7–9 |
| SCAN | Multiple ADRCs | GE; Philips; | 3 | MPRAGE; FSPGR; TFE | 1 × 1 × 1 | ~6.5† or 2300† | 2.82–3.12 | 400–900 | 9–11 |
| Temple Social | Single site | Siemens | 3 | MPRAGE | 1 × 1 × 1 | 2400 | 2.17–2.18 | 1150 | 8 |
| UCLA LA5c | 2 UCLA sites | Siemens | 3 | MPRAGE | 1 × 1 × 1 | 2530 | 3.31–3.52 | 1100 | 7 |
| UCSD ACE | Toddler cohort | GE | 1.5 | IR-FSPGR | 0.9375 × 0.9375 × 1.2 | 6.496–6.796† | 2.796–2.928 | 600 | 12 |
| VMAP 2.0 | VMAP 2.0 | Philips | 3 | MPRAGE | 1 × 1 × 1 | 6.33–6.57† | 2.924–2.979 | NR | 9 |
| VMAP/TAP | VMAP / Tennessee Alzheimer's Project | Philips | 3 | MPRAGE | 1 × 1 × 1 | 6.44–6.57† | 2.929–2.948 | NR | 9 |
| Washington University / Knight ADRC | Knight ADRC; multiple scanner groups | Siemens | 3 | MPRAGE | Trio 1 × 1 × 1; mMR 1 × 1 × 1.2 | Trio 2400; mMR 2300 | Trio 3.16; mMR 2.95 | Trio 1000; mMR 900 | Trio 8; mMR 9 |
| WRAP | Multiple GE scanner generations | GE | 3 | IR-SPGR; BRAVO | 1 × 1 × 1 | 6.52–8.16† | 2.816–3.18 | 400–900 | 8–12 |

*NR, not reported; NA, not applicable.*

*Voxel dimensions denote reconstructed image spacing.*

*† Anatomical TR may represent excitation TR (~6–9 ms) or preparation-cycle TR (~1900–3000 ms), depending on vendor and sequence; values using different definitions should not be averaged.*

*Protocol choices and ranges summarize published cohort protocols and bounded, read-only cohort-level BIDS sampling. The BIDS sample was not joined scan-by-scan to the final manifest; see the detailed acquisition workbook for sources and dataset-specific caveats.*

***Supplementary Data File 1****. Attached CSVs with sex-stratified percentiles for all morphometric features, at all cervical levels.*

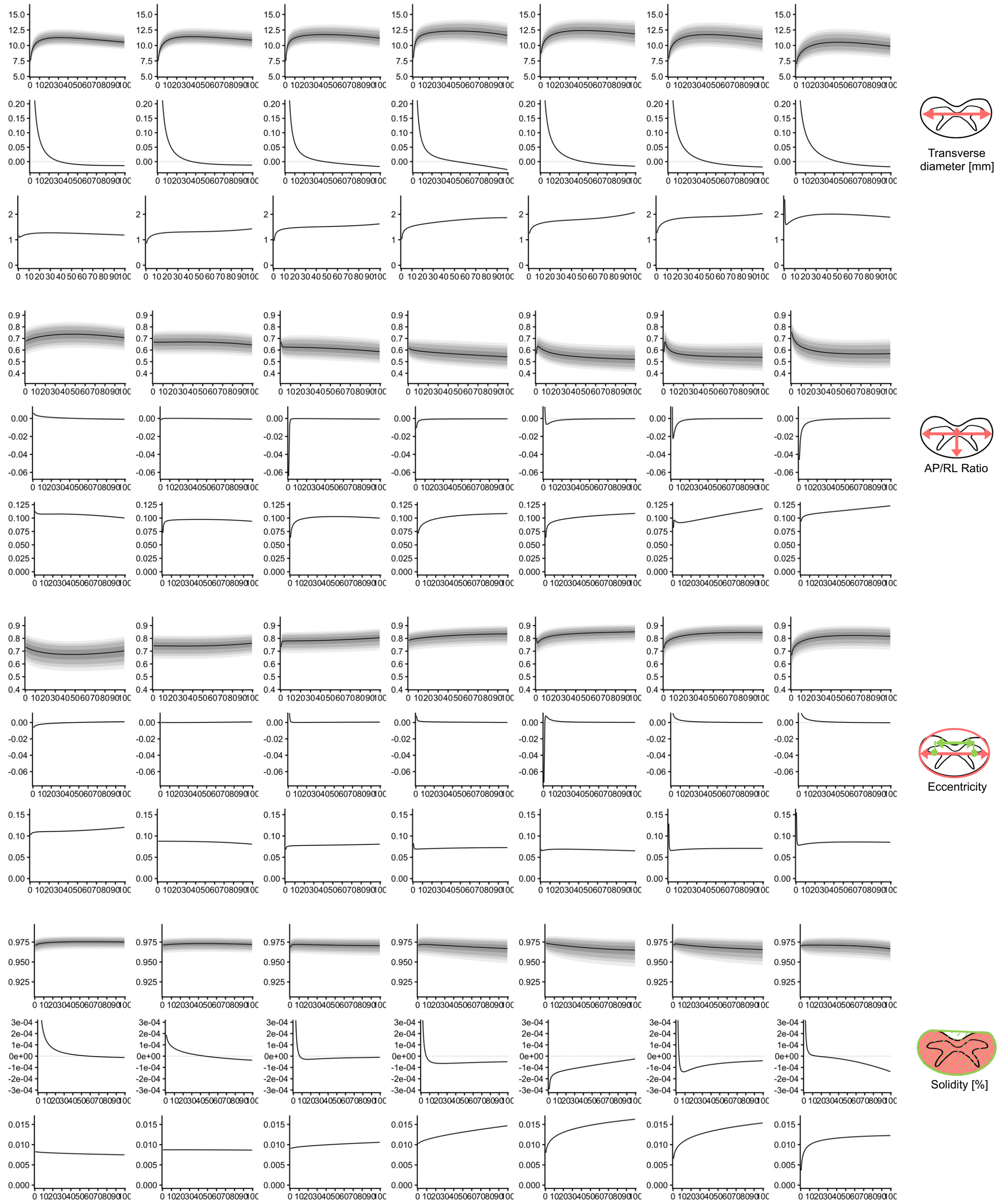

***Supplementary Figure 3**. Lifespan Dynamics of Spinal Cord Morphometry Across the Cervical Spine. Figure displays the normative trajectory (centiles), annualized rate of change (change-per-year), and normative population variability (standard deviation, σ) for Transverse Diameter, Compression Ratio, Eccentricity, Solidity. Normative trajectories reveal distinct developmental and aging patterns that vary by vertebral level; annualized rate of change quantifies these dynamics, highlighting periods of rapid growth in childhood and adolescence followed by a steady rate of atrophy in late adulthood; Population variability illustrates the spread of measurements changes across the lifespan.*

***Supplementary Table 3**. Quantifying Annualized rates of change for clinical benchmarking. Change Per Year and Percent Change Per Year are given for all features (at all levels), for each lifespan stage. Brain measurements are also given for reference.*

| Measure | Vertebral Level | Change Per Year | | | | | | | Percent Change Per Year | | | | | | |
|---|---|---|---|---|---|---|---|---|---|---|---|---|---|---|---|
| | | Infancy | Early Childhood | Late Childhood | Adolescence | Young Adulthood | Middle Adulthood | Late Adulthood | Infancy | Early Childhood | Late Childhood | Adolescence | Young Adulthood | Middle Adulthood | Late Adulthood |
| CSA | C1 | 7.9 | 3.5 | 1.4 | 0.67 | 0.16 | -0.12 | -0.24 | 21 | 7.4 | 2.3 | 0.99 | 0.22 | -0.17 | -0.37 |
| | C2 | 10 | 3.8 | 1.2 | 0.54 | 0.13 | -0.087 | -0.19 | 31 | 8.3 | 2 | 0.83 | 0.19 | -0.13 | -0.29 |
| | C3 | 9.7 | 3.3 | 0.98 | 0.43 | 0.083 | -0.094 | -0.17 | 26 | 6.9 | 1.6 | 0.66 | 0.12 | -0.14 | -0.28 |
| | C4 | 18 | 4.5 | 0.77 | 0.29 | 0.044 | -0.094 | -0.2 | 54 | 9.1 | 1.2 | 0.43 | 0.065 | -0.14 | -0.32 |
| | C5 | 17 | 4.1 | 0.69 | 0.26 | 0.038 | -0.088 | -0.19 | 50 | 8.4 | 1.1 | 0.4 | 0.058 | -0.14 | -0.32 |
| | C6 | 20 | 4 | 0.64 | 0.26 | 0.079 | -0.04 | -0.19 | 75 | 9.5 | 1.2 | 0.47 | 0.14 | -0.068 | -0.35 |
| | C7 | 8 | 2.4 | 0.67 | 0.3 | 0.067 | -0.052 | -0.11 | 29 | 6.7 | 1.5 | 0.63 | 0.14 | -0.11 | -0.23 |
| Diameter AP | C1 | 0.68 | 0.28 | 0.1 | 0.047 | 0.011 | -0.008 | -0.017 | 12 | 4.4 | 1.4 | 0.6 | 0.14 | -0.098 | -0.22 |
| | C2 | 0.53 | 0.21 | 0.073 | 0.033 | 0.0075 | -0.0061 | -0.012 | 9.5 | 3.4 | 1 | 0.46 | 0.099 | -0.081 | -0.17 |
| | C3 | 0.47 | 0.18 | 0.059 | 0.025 | 0.0033 | -0.0077 | -0.012 | 8.3 | 3 | 0.86 | 0.35 | 0.045 | -0.11 | -0.18 |
| | C4 | 0.58 | 0.18 | 0.043 | 0.015 | -0.0012 | -0.009 | -0.012 | 10 | 2.9 | 0.62 | 0.21 | -0.017 | -0.13 | -0.19 |
| | C5 | 0.84 | 0.17 | 0.024 | 0.0071 | -0.0018 | -0.0077 | -0.013 | 15 | 2.8 | 0.35 | 0.1 | -0.026 | -0.12 | -0.21 |
| | C6 | 0.71 | 0.15 | 0.021 | 0.0082 | 0.0014 | -0.0042 | -0.013 | 14 | 2.5 | 0.34 | 0.13 | 0.022 | -0.065 | -0.22 |
| | C7 | 0.2 | 0.063 | 0.019 | 0.0098 | 0.0037 | -0.0019 | -0.011 | 3.8 | 1.1 | 0.32 | 0.17 | 0.063 | -0.031 | -0.19 |
| Diameter RL | C1 | 1.5 | 0.46 | 0.1 | 0.036 | 0.0034 | -0.0093 | -0.013 | 20 | 5.1 | 0.97 | 0.33 | 0.031 | -0.084 | -0.12 |
| | C2 | 1.5 | 0.47 | 0.11 | 0.042 | 0.0078 | -0.0062 | -0.011 | 21 | 5.2 | 1 | 0.38 | 0.069 | -0.054 | -0.099 |
| | C3 | 2.4 | 0.54 | 0.084 | 0.032 | 0.0085 | -0.0036 | -0.012 | 32 | 5.7 | 0.76 | 0.28 | 0.072 | -0.031 | -0.11 |
| | C4 | 2.4 | 0.54 | 0.085 | 0.034 | 0.011 | -0.0017 | -0.017 | 30 | 5.4 | 0.74 | 0.29 | 0.093 | -0.014 | -0.14 |
| | C5 | 1.2 | 0.39 | 0.11 | 0.051 | 0.015 | -0.0027 | -0.012 | 14 | 3.9 | 0.97 | 0.43 | 0.13 | -0.022 | -0.1 |
| | C6 | 1.3 | 0.41 | 0.12 | 0.053 | 0.014 | -0.0052 | -0.015 | 16 | 4.5 | 1.1 | 0.47 | 0.12 | -0.045 | -0.13 |
| | C7 | 0.93 | 0.33 | 0.12 | 0.06 | 0.019 | -0.0031 | -0.014 | 13 | 4.1 | 1.3 | 0.6 | 0.19 | -0.03 | -0.14 |
| Compression Ratio | C1 | 0.006 | 0.0038 | 0.0023 | 0.0015 | 0.00068 | -0.000055 | -0.0007 | 0.88 | 0.56 | 0.33 | 0.21 | 0.093 | -0.0075 | -0.097 |
| | C2 | -0.0018 | -0.00022 | 0.000084 | 0.0001 | 0.000038 | -0.00015 | -0.00056 | -0.26 | -0.033 | 0.013 | 0.015 | 0.0057 | -0.023 | -0.085 |
| | C3 | -0.064 | -0.0055 | -0.00017 | -0.00016 | -0.00025 | -0.00039 | -0.00061 | -9.6 | -0.86 | -0.027 | -0.025 | -0.04 | -0.064 | -0.1 |
| | C4 | -0.011 | -0.0036 | -0.0013 | -0.00089 | -0.00065 | -0.00053 | -0.00047 | -1.7 | -0.59 | -0.22 | -0.15 | -0.11 | -0.094 | -0.085 |
| | C5 | 0.1 | 0.003 | -0.0041 | -0.0023 | -0.0011 | -0.00058 | -0.00033 | 18 | 0.48 | -0.68 | -0.39 | -0.2 | -0.11 | -0.062 |
| | C6 | 0.17 | -0.0047 | -0.0046 | -0.0018 | -0.00064 | -0.00026 | -0.00013 | 30 | -0.81 | -0.79 | -0.31 | -0.12 | -0.048 | -0.024 |
| | C7 | -0.046 | -0.017 | -0.0056 | -0.0028 | -0.0012 | -0.00039 | 0.000041 | -6.1 | -2.5 | -0.87 | -0.46 | -0.21 | -0.068 | 0.0072 |
| Eccentricity | C1 | -0.0064 | -0.004 | -0.0023 | -0.0014 | -0.00061 | 0.000061 | 0.00066 | -0.87 | -0.55 | -0.32 | -0.21 | -0.089 | 0.009 | 0.096 |
| | C2 | -0.000016 | -0.000041 | -0.000064 | -0.000062 | -0.0000033 | 0.00014 | 0.00047 | -0.0022 | -0.0055 | -0.0086 | -0.0083 | -0.00045 | 0.019 | 0.062 |
| | C3 | 0.07 | 0.0058 | 0.000077 | 0.00008 | 0.00015 | 0.00025 | 0.00041 | 9.5 | 0.75 | 0.0098 | 0.01 | 0.019 | 0.032 | 0.052 |
| | C4 | 0.0082 | 0.0023 | 0.0011 | 0.00081 | 0.00058 | 0.00039 | 0.00018 | 1 | 0.29 | 0.14 | 0.1 | 0.071 | 0.048 | 0.021 |
| | C5 | -0.073 | 0.0015 | 0.0031 | 0.0015 | 0.00069 | 0.00034 | 0.00018 | -9.1 | 0.19 | 0.39 | 0.19 | 0.083 | 0.04 | 0.022 |
| | C6 | 0.048 | 0.01 | 0.0032 | 0.0018 | 0.00084 | 0.00032 | 0.000019 | 6.7 | 1.4 | 0.41 | 0.22 | 0.1 | 0.037 | 0.0023 |
| | C7 | 0.065 | 0.014 | 0.0043 | 0.0023 | 0.00095 | 0.00022 | -0.00018 | 9.8 | 2 | 0.56 | 0.29 | 0.12 | 0.027 | -0.022 |
| Solidity | C1 | 0.0012 | 0.00046 | 0.00016 | 0.000079 | 0.000031 | 0.0000058 | -0.0000076 | 0.12 | 0.047 | 0.016 | 0.0081 | 0.0032 | 0.0006 | -0.00078 |
| | C2 | 0.00019 | 0.00013 | 0.000076 | 0.000049 | 0.00002 | -0.0000042 | -0.000026 | 0.019 | 0.013 | 0.0078 | 0.005 | 0.002 | -0.00043 | -0.0026 |
| | C3 | 0.0041 | 0.00058 | -0.0000082 | -0.000027 | -0.000024 | -0.000018 | -0.000013 | 0.42 | 0.06 | -0.00085 | -0.0028 | -0.0024 | -0.0018 | -0.0013 |
| | C4 | 0.0013 | 0.00025 | -0.000032 | -0.00006 | -0.000064 | -0.000059 | -0.000053 | 0.14 | 0.026 | -0.0033 | -0.0062 | -0.0066 | -0.0061 | -0.0054 |
| | C5 | -0.00029 | -0.00019 | -0.00015 | -0.00013 | -0.00011 | -0.000083 | -0.000048 | -0.03 | -0.02 | -0.015 | -0.014 | -0.011 | -0.0086 | -0.0049 |
| | C6 | 0.0021 | 0.00019 | -0.00014 | -0.00012 | -0.00009 | -0.000066 | -0.000048 | 0.22 | 0.02 | -0.014 | -0.012 | -0.0092 | -0.0068 | -0.005 |
| | C7 | 0.0013 | 0.0002 | 0.000012 | -2.9E-07 | -0.000012 | -0.000035 | -0.00009 | 0.13 | 0.021 | 0.0013 | -0.00003 | -0.0012 | -0.0036 | -0.0093 |
| Length | C1 | 6.7 | 1.4 | 0.37 | 0.17 | 0.046 | -0.0084 | -0.028 | 200 | 17 | 2.8 | 1.1 | 0.28 | -0.05 | -0.18 |
| | C2 | 7.4 | 1.4 | 0.34 | 0.15 | 0.043 | -0.0064 | -0.025 | 250 | 17 | 2.6 | 1 | 0.27 | -0.04 | -0.16 |
| | C3 | 9.3 | 1.4 | 0.41 | 0.24 | 0.088 | -0.0047 | -0.052 | 3100 | 23 | 3.8 | 1.8 | 0.6 | -0.03 | -0.35 |
| | C4 | 2.9 | 0.95 | 0.42 | 0.25 | 0.097 | -0.0011 | -0.052 | 80 | 15 | 4.1 | 2 | 0.67 | -0.0069 | -0.36 |
| | C5 | 7.4 | 1.3 | 0.46 | 0.27 | 0.11 | -0.00071 | -0.057 | 1100 | 23 | 4.5 | 2.1 | 0.73 | -0.0044 | -0.39 |
| | C6 | 12 | 1.4 | 0.32 | 0.18 | 0.061 | -0.0085 | -0.042 | 3400 | 19 | 2.9 | 1.4 | 0.43 | -0.058 | -0.31 |
| | C7 | -12 | -0.39 | 0.22 | 0.1 | 0.041 | 0.017 | 0.008 | -81 | -4.6 | 2.2 | 0.89 | 0.34 | 0.14 | 0.061 |
| Brain-Stem | | 4200 | 1600 | 470 | 190 | 39 | -28 | -49 | 56 | 13 | 2.6 | 0.96 | 0.18 | -0.13 | -0.24 |
| BrainSegVol | | 430000 | 94000 | 8200 | -450 | -2900 | -3000 | -2600 | 71 | 10 | 0.69 | -0.037 | -0.24 | -0.27 | -0.24 |
| BrainSegVolNotVent | | 370000 | 77000 | 8400 | 480 | -2700 | -3600 | -3700 | 54 | 8 | 0.71 | 0.04 | -0.23 | -0.33 | -0.37 |
| CerebralWhiteMatter Vol | | 100000 | 30000 | 8700 | 3800 | 550 | -1100 | -1700 | 48 | 9.8 | 2.2 | 0.84 | 0.12 | -0.24 | -0.42 |
| CortexVol | | 300000 | 28000 | 260 | -61 | -370 | -1000 | -2300 | 110 | 6.6 | 0.053 | -0.013 | -0.077 | -0.21 | -0.56 |
| eTIV | | -440000 | 120000 | 12000 | 2400 | 490 | 100 | 29 | -48 | 11 | 0.85 | 0.17 | 0.033 | 0.007 | 0.002 |
| SubCortGrayVol | | 19000 | 4700 | 600 | 78 | -140 | -210 | -210 | 66 | 10 | 1 | 0.13 | -0.23 | -0.36 | -0.42 |
| TotalGrayVol | | 400000 | 61000 | -2100 | -3900 | -3300 | -2500 | -1800 | 120 | 10 | -0.28 | -0.53 | -0.48 | -0.39 | -0.32 |
| TotalThalamusVol | | 3500 | 1100 | 210 | 29 | -50 | -68 | -62 | 44 | 9.6 | 1.4 | 0.18 | -0.32 | -0.48 | -0.5 |

*Supplementary Table 4. Population variability across the lifespan. Coefficient of variation, and population standard deviation are given for all features (at all levels), for each lifespan stage. Brain measurements are also given for reference.*

| Measure | Vertebral Level | Population Standard Deviation | | | | | | | Coefficient of Variation | | | | | | |
|---|---|---|---|---|---|---|---|---|---|---|---|---|---|---|---|
| | | Infancy | Early Childhood | Late Childhood | Adolescence | Young Adulthood | Middle Adulthood | Late Adulthood | Infancy | Early Childhood | Late Childhood | Adolescence | Young Adulthood | Middle Adulthood | Late Adulthood |
| CSA | C1 | 3.2 | 4.3 | 5.3 | 5.9 | 6.4 | 6.5 | 6.3 | 8.6 | 8.6 | 8.6 | 8.7 | 8.7 | 8.9 | 9.4 |
| | C2 | 2.8 | 4.1 | 5 | 5.4 | 5.8 | 6 | 6.1 | 8.4 | 8.4 | 8.4 | 8.4 | 8.5 | 8.8 | 9.6 |
| | C3 | 3.4 | 4.7 | 5.5 | 5.9 | 6.3 | 6.4 | 6.5 | 9.1 | 9.2 | 9.2 | 9.2 | 9.3 | 9.6 | 10 |
| | C4 | 3.3 | 5.4 | 6.3 | 6.6 | 6.9 | 7.1 | 7.3 | 9.9 | 9.9 | 9.9 | 10 | 10 | 11 | 12 |
| | C5 | 3.7 | 5.8 | 6.7 | 7 | 7.2 | 7.3 | 7.5 | 11 | 11 | 11 | 11 | 11 | 11 | 12 |
| | C6 | 10 | 6.2 | 6.8 | 7.1 | 7.4 | 7.4 | 6.9 | 39 | 13 | 13 | 13 | 13 | 13 | 13 |
| | C7 | 7.8 | 5.7 | 6.5 | 6.9 | 7.2 | 7.2 | 6.8 | 28 | 15 | 15 | 15 | 15 | 15 | 15 |
| Diameter AP | C1 | 0.33 | 0.39 | 0.44 | 0.47 | 0.49 | 0.5 | 0.5 | 6 | 6 | 6 | 6 | 6 | 6.1 | 6.4 |
| | C2 | 0.29 | 0.34 | 0.37 | 0.4 | 0.42 | 0.44 | 0.45 | 5.2 | 5.3 | 5.3 | 5.4 | 5.6 | 5.8 | 6.2 |
| | C3 | 0.3 | 0.34 | 0.39 | 0.41 | 0.44 | 0.46 | 0.47 | 5.3 | 5.4 | 5.6 | 5.8 | 6.1 | 6.4 | 6.8 |
| | C4 | 0.35 | 0.4 | 0.43 | 0.45 | 0.47 | 0.49 | 0.5 | 6.2 | 6.2 | 6.3 | 6.4 | 6.7 | 7 | 7.6 |
| | C5 | 0.34 | 0.41 | 0.45 | 0.47 | 0.49 | 0.51 | 0.52 | 6.2 | 6.4 | 6.7 | 6.9 | 7.3 | 7.7 | 8.2 |
| | C6 | 0.38 | 0.44 | 0.46 | 0.46 | 0.47 | 0.48 | 0.53 | 7.3 | 7.3 | 7.3 | 7.3 | 7.3 | 7.6 | 8.7 |
| | C7 | 0.37 | 0.41 | 0.43 | 0.45 | 0.48 | 0.5 | 0.52 | 7 | 7.2 | 7.5 | 7.7 | 8.1 | 8.5 | 9 |
| Diameter RL | C1 | 0.58 | 0.57 | 0.61 | 0.63 | 0.64 | 0.63 | 0.61 | 7.7 | 6 | 5.7 | 5.7 | 5.6 | 5.6 | 5.6 |
| | C2 | 0.43 | 0.54 | 0.61 | 0.63 | 0.65 | 0.66 | 0.68 | 5.7 | 5.7 | 5.7 | 5.7 | 5.7 | 5.8 | 6.2 |
| | C3 | 0.48 | 0.64 | 0.71 | 0.73 | 0.75 | 0.76 | 0.78 | 6.4 | 6.4 | 6.4 | 6.4 | 6.4 | 6.5 | 6.8 |
| | C4 | 0.51 | 0.68 | 0.75 | 0.79 | 0.83 | 0.88 | 0.92 | 6.4 | 6.4 | 6.5 | 6.6 | 6.8 | 7.1 | 7.7 |
| | C5 | 0.62 | 0.73 | 0.81 | 0.84 | 0.87 | 0.9 | 0.96 | 7.1 | 7.1 | 7.1 | 7.1 | 7.1 | 7.3 | 7.9 |
| | C6 | 0.63 | 0.77 | 0.86 | 0.9 | 0.93 | 0.95 | 0.98 | 8 | 8 | 8 | 8 | 8 | 8.1 | 8.6 |
| | C7 | 1.3 | 0.82 | 0.89 | 0.95 | 0.99 | 1 | 0.97 | 18 | 9.9 | 9.6 | 9.5 | 9.5 | 9.5 | 9.5 |
| Compression Ratio | C1 | 0.056 | 0.054 | 0.054 | 0.054 | 0.054 | 0.053 | 0.052 | 8.3 | 7.9 | 7.6 | 7.5 | 7.4 | 7.2 | 7.1 |
| | C2 | 0.036 | 0.046 | 0.048 | 0.048 | 0.049 | 0.049 | 0.048 | 5.4 | 6.9 | 7.2 | 7.2 | 7.3 | 7.3 | 7.3 |
| | C3 | 0.032 | 0.043 | 0.048 | 0.05 | 0.051 | 0.051 | 0.051 | 4.8 | 6.8 | 7.6 | 8 | 8.2 | 8.4 | 8.5 |
| | C4 | 0.036 | 0.042 | 0.046 | 0.048 | 0.051 | 0.053 | 0.054 | 5.7 | 6.9 | 7.7 | 8.2 | 8.8 | 9.3 | 9.8 |
| | C5 | 0.032 | 0.043 | 0.047 | 0.048 | 0.05 | 0.052 | 0.053 | 5.6 | 6.9 | 7.8 | 8.4 | 9 | 9.6 | 10 |
| | C6 | 0.041 | 0.047 | 0.046 | 0.046 | 0.049 | 0.052 | 0.056 | 7 | 7.4 | 7.8 | 8.2 | 8.8 | 9.5 | 10 |
| | C7 | 0.047 | 0.051 | 0.053 | 0.054 | 0.055 | 0.057 | 0.06 | 6.2 | 7.5 | 8.3 | 8.9 | 9.5 | 10 | 11 |
| Eccentricity | C1 | 0.051 | 0.054 | 0.055 | 0.055 | 0.055 | 0.056 | 0.058 | 7 | 7.5 | 7.8 | 8 | 8.2 | 8.3 | 8.5 |
| | C2 | 0.044 | 0.044 | 0.044 | 0.044 | 0.044 | 0.043 | 0.042 | 5.9 | 5.9 | 5.9 | 5.9 | 5.9 | 5.8 | 5.6 |
| | C3 | 0.034 | 0.038 | 0.039 | 0.039 | 0.039 | 0.039 | 0.04 | 4.6 | 4.9 | 4.9 | 5 | 5 | 5 | 5 |
| | C4 | 0.041 | 0.035 | 0.035 | 0.035 | 0.036 | 0.036 | 0.036 | 5.3 | 4.4 | 4.4 | 4.4 | 4.4 | 4.4 | 4.4 |
| | C5 | 0.034 | 0.033 | 0.034 | 0.034 | 0.034 | 0.034 | 0.033 | 4.3 | 4.3 | 4.2 | 4.2 | 4.1 | 4.1 | 3.9 |
| | C6 | 0.065 | 0.034 | 0.034 | 0.034 | 0.035 | 0.035 | 0.035 | 8.9 | 4.4 | 4.2 | 4.2 | 4.2 | 4.2 | 4.2 |
| | C7 | 0.078 | 0.04 | 0.04 | 0.041 | 0.042 | 0.043 | 0.043 | 12 | 5.5 | 5.2 | 5.2 | 5.2 | 5.2 | 5.2 |
| Solidity | C1 | 0.0041 | 0.0041 | 0.004 | 0.004 | 0.0039 | 0.0039 | 0.0038 | 0.43 | 0.42 | 0.42 | 0.41 | 0.4 | 0.4 | 0.39 |
| | C2 | 0.0044 | 0.0044 | 0.0044 | 0.0044 | 0.0044 | 0.0044 | 0.0043 | 0.45 | 0.45 | 0.45 | 0.45 | 0.45 | 0.45 | 0.45 |
| | C3 | 0.0045 | 0.0046 | 0.0047 | 0.0048 | 0.0049 | 0.005 | 0.0052 | 0.47 | 0.47 | 0.48 | 0.49 | 0.51 | 0.52 | 0.54 |
| | C4 | 0.0051 | 0.0053 | 0.0056 | 0.0058 | 0.0061 | 0.0065 | 0.007 | 0.53 | 0.55 | 0.57 | 0.6 | 0.63 | 0.67 | 0.73 |
| | C5 | 0.004 | 0.005 | 0.0058 | 0.0063 | 0.0069 | 0.0074 | 0.0079 | 0.41 | 0.52 | 0.6 | 0.65 | 0.71 | 0.76 | 0.82 |
| | C6 | 0.0033 | 0.0043 | 0.0051 | 0.0057 | 0.0063 | 0.0068 | 0.0074 | 0.34 | 0.44 | 0.53 | 0.58 | 0.65 | 0.71 | 0.76 |
| | C7 | 0.0018 | 0.0038 | 0.0048 | 0.0053 | 0.0057 | 0.0059 | 0.0061 | 0.19 | 0.39 | 0.5 | 0.54 | 0.58 | 0.61 | 0.63 |
| Length | C1 | 3.4 | 1.7 | 1.8 | 1.9 | 2 | 2 | 1.9 | 100 | 18 | 13 | 12 | 12 | 12 | 12 |
| | C2 | 3 | 1.4 | 1.4 | 1.5 | 1.5 | 1.5 | 1.5 | 100 | 14 | 10 | 9.9 | 9.6 | 9.5 | 9.4 |
| | C3 | 2.5 | 1.8 | 1.7 | 1.9 | 2.2 | 2.2 | 2 | 820 | 25 | 16 | 15 | 14 | 14 | 14 |
| | C4 | 2.1 | 1.4 | 1.8 | 2.1 | 2.4 | 2.5 | 2.3 | 59 | 20 | 17 | 16 | 16 | 16 | 16 |
| | C5 | 2.6 | 1.8 | 1.9 | 2.2 | 2.5 | 2.6 | 2.4 | 410 | 27 | 18 | 17 | 16 | 16 | 16 |
| | C6 | 2.5 | 1.7 | 1.9 | 2.2 | 2.4 | 2.5 | 2.4 | 700 | 20 | 17 | 17 | 17 | 17 | 17 |
| | C7 | 12 | 1.8 | 1.7 | 1.8 | 1.8 | 1.9 | 1.9 | 80 | 21 | 16 | 15 | 15 | 15 | 15 |
| Brain-Stem | | 730 | 1300 | 1700 | 1900 | 2100 | 2100 | 2100 | 9.5 | 9.5 | 9.5 | 9.6 | 9.6 | 9.8 | 10 |
| BrainSegVol | | 100000 | 90000 | 100000 | 100000 | 98000 | 93000 | 86000 | 17 | 8.6 | 8.3 | 8.3 | 8.3 | 8.3 | 8.3 |
| BrainSegVolNotVent | | 150000 | 90000 | 97000 | 98000 | 96000 | 91000 | 82000 | 23 | 8.7 | 8.2 | 8.2 | 8.2 | 8.2 | 8.2 |
| CerebralWhiteMatter Vol | | 93000 | 38000 | 44000 | 48000 | 51000 | 50000 | 45000 | 45 | 12 | 11 | 11 | 11 | 11 | 11 |
| CortexVol | | 23000 | 40000 | 41000 | 41000 | 41000 | 41000 | 38000 | 8.5 | 8.5 | 8.5 | 8.5 | 8.6 | 8.7 | 9.2 |
| eTIV | | 140000 | 120000 | 140000 | 140000 | 140000 | 140000 | 140000 | 15 | 9.9 | 9.7 | 9.7 | 9.7 | 9.7 | 9.7 |
| SubCortGrayVol | | 2100 | 3600 | 4200 | 4300 | 4300 | 4200 | 4200 | 7 | 7 | 7 | 7 | 7.1 | 7.3 | 8.3 |
| TotalGrayVol | | 26000 | 52000 | 55000 | 54000 | 50000 | 47000 | 44000 | 7.4 | 7.4 | 7.4 | 7.4 | 7.4 | 7.5 | 7.9 |
| TotalThalamusVol | | 500 | 930 | 1200 | 1400 | 1400 | 1400 | 1200 | 6.1 | 7.2 | 8 | 8.4 | 9 | 9.4 | 9.9 |

### Out of Sample Alignment validation: (VUIIS-MS)

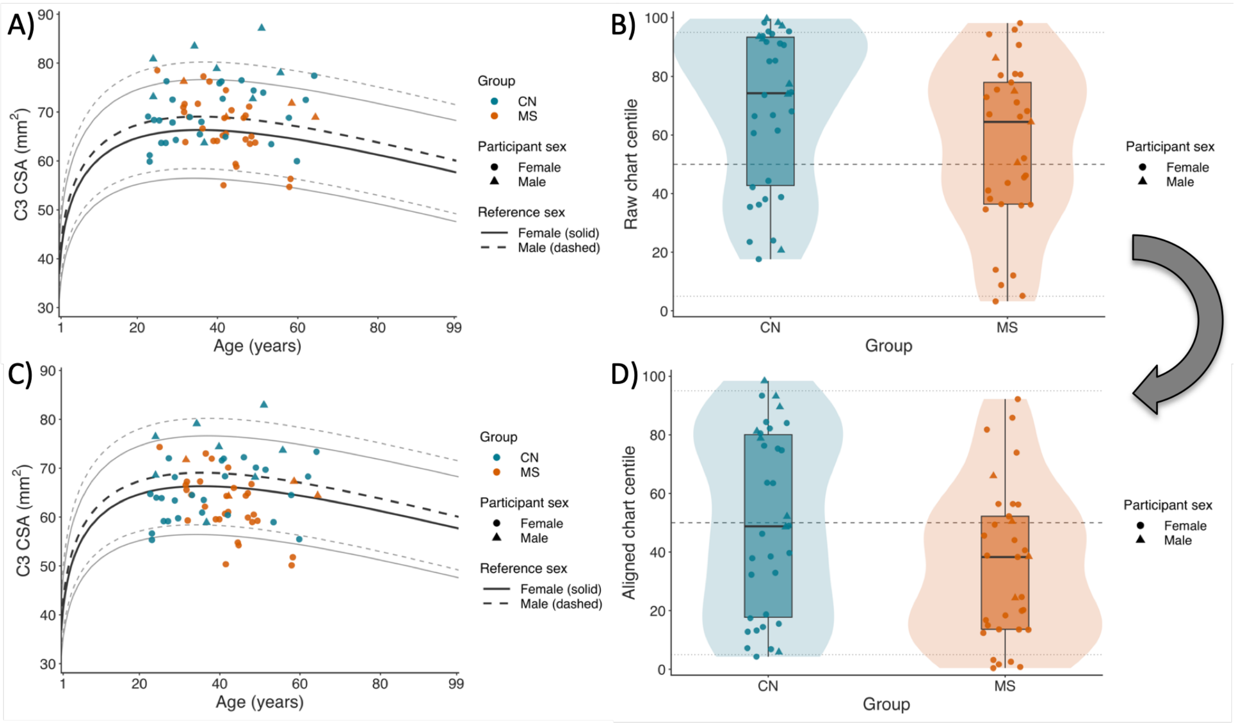


***Supplementary Figure 4****. Out-of-sample alignment following an introduced 5% measurement bias. The analysis and panel organization match Figure 8, but every C3 CSA measurement was multiplied by 1.05 before alignment. (A) Biased C3 CSA plotted against the age- and sex-specific reference chart. (B) Distributions of raw chart centiles after introducing the bias. (C) Harmonized C3 CSA after offsets estimated from controls were applied to both groups. (D) Distributions of aligned chart centiles. Alignment recovered the imposed scale factor to numerical precision: the estimated change in the location offset was 0.04879016, matching log(1.05)=0.04879016, and aligned centiles differed from the original unshifted analysis by at most 5.2x10^-6 centile points. Reference curves and line styles follow Figure 8.*

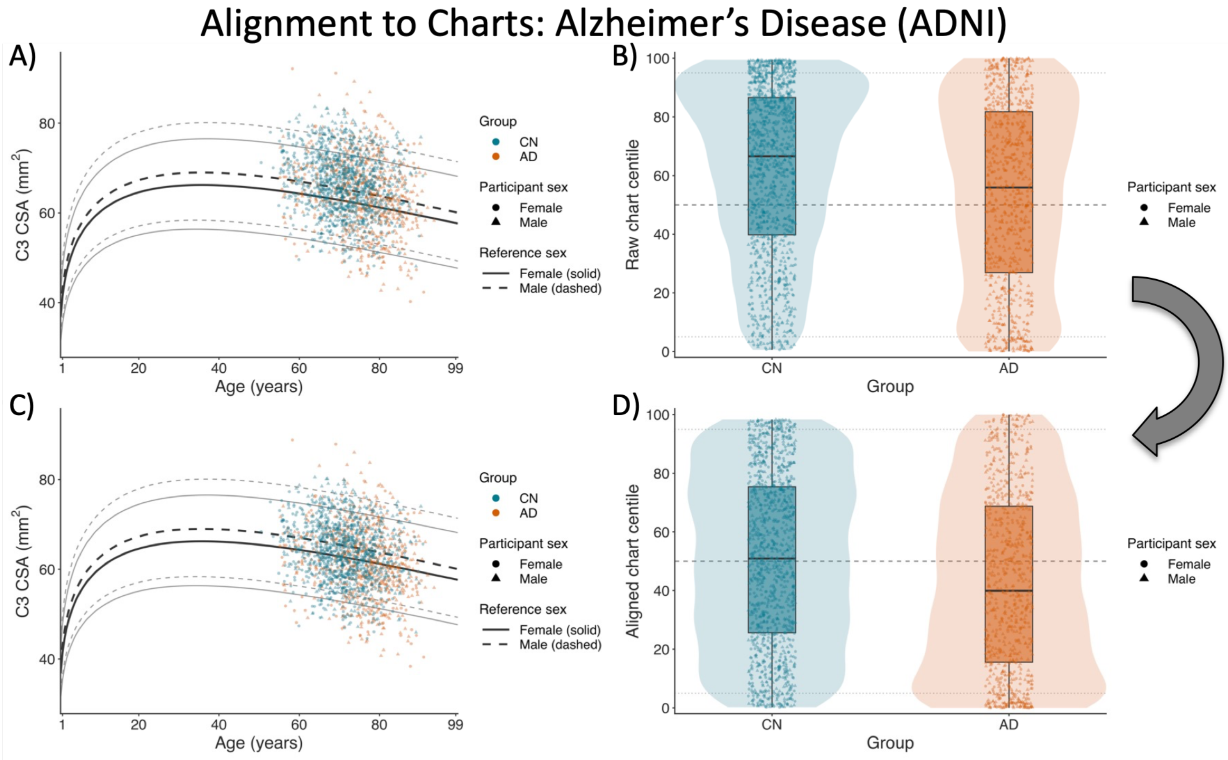


***Supplementary Figure 5.*** *Out-of-sample alignment in the Alzheimer's Disease Neuroimaging Initiative. The example included 1,157 controls and 748 participants with Alzheimer disease and was not used to build the C3 reference chart used for this analysis. (A) Observed C3 CSA plotted against the age- and sex-specific reference chart. (B) Distributions of raw chart centiles in controls and participants with Alzheimer disease. (C) Harmonized C3 CSA after offsets estimated from controls were applied unchanged to both groups. (D) Distributions of aligned chart centiles. Following alignment, mean and median centiles were 50.5 and 50.9 in controls and 43.0 and 39.9 in Alzheimer disease; 13.8% of participants with Alzheimer disease and 6.1% of controls were below the 5th centile. Reference curves and line styles follow Figure 8.*

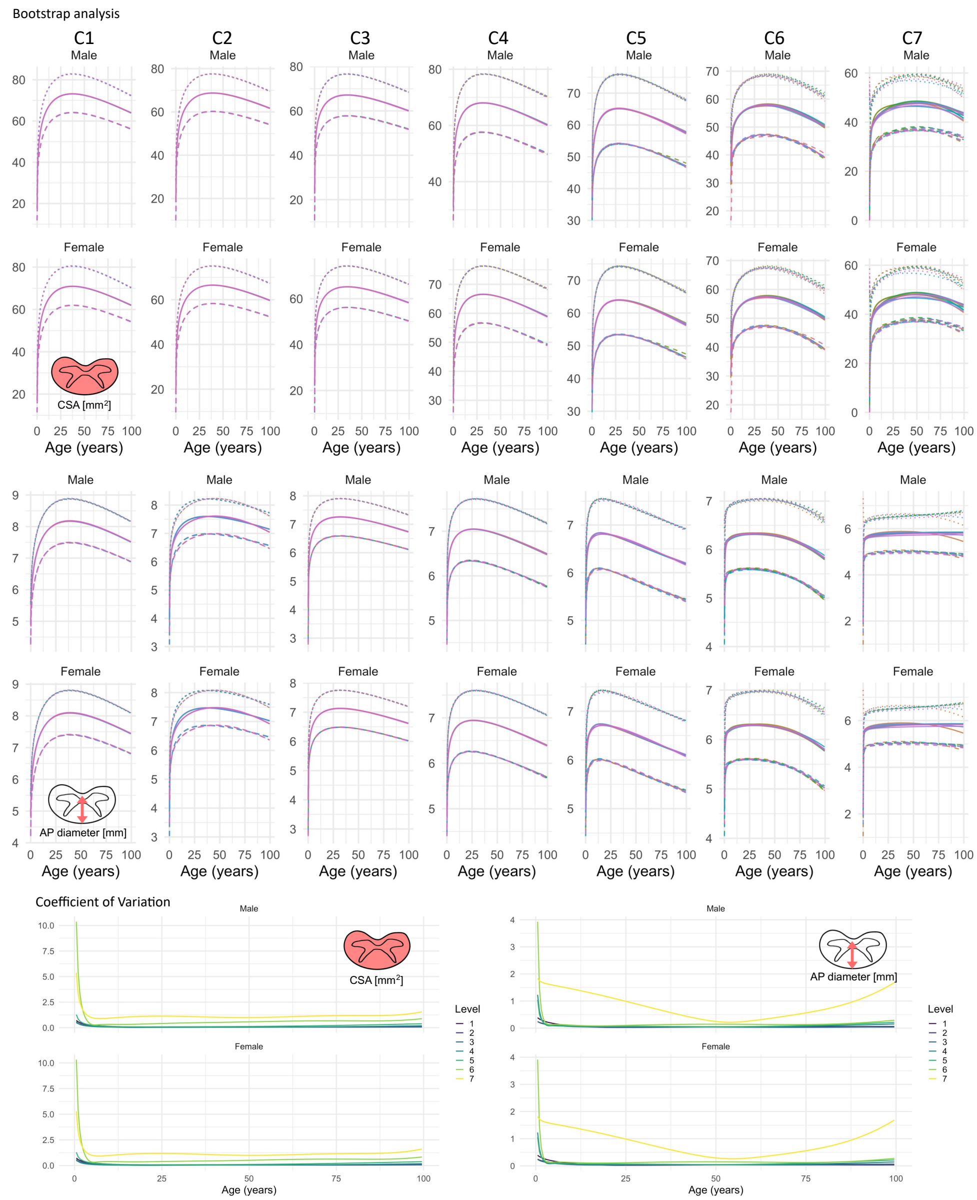


***Supplementary Figure 6.*** *Stability analysis. Cross-sectional datasets were randomly selected and a bootstrapping procedure was used to quantify uncertainty in models. Centiles are shown for CSA and AP diameter, across all cervical levels, with coefficient of variation (CV) <1% for all features and levels with the exception of C7 (and in infants).*

Trajectories for each left-out dataset, visual check of stability and age windows with greater divergence

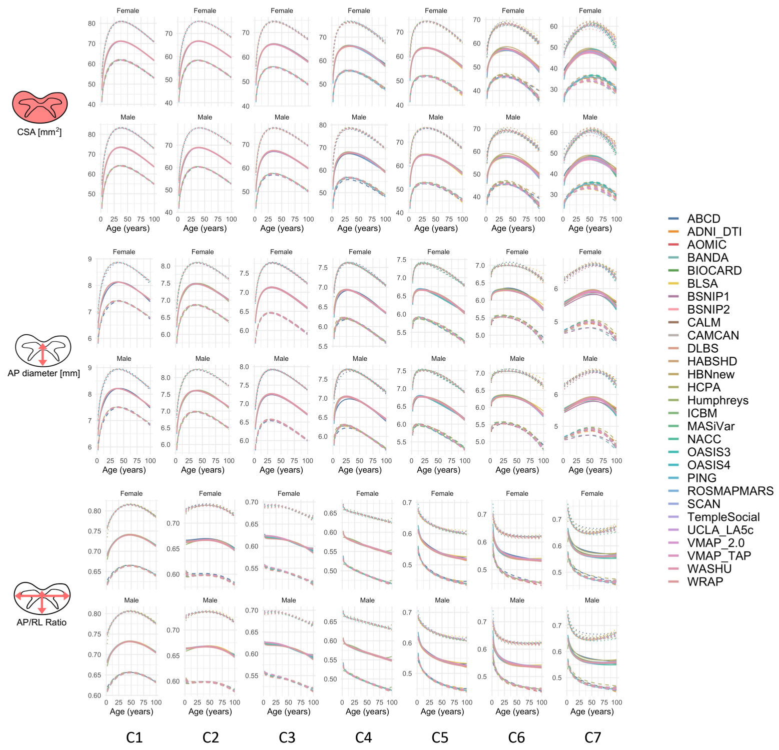


***Supplementary Figure 7.*** *Leave-one-out site analysis to assess robustness of models to specific sites. Visually, centiles scores remain similar, with more site-driven variation at C7. Left-out datasets are shown as different colors.*

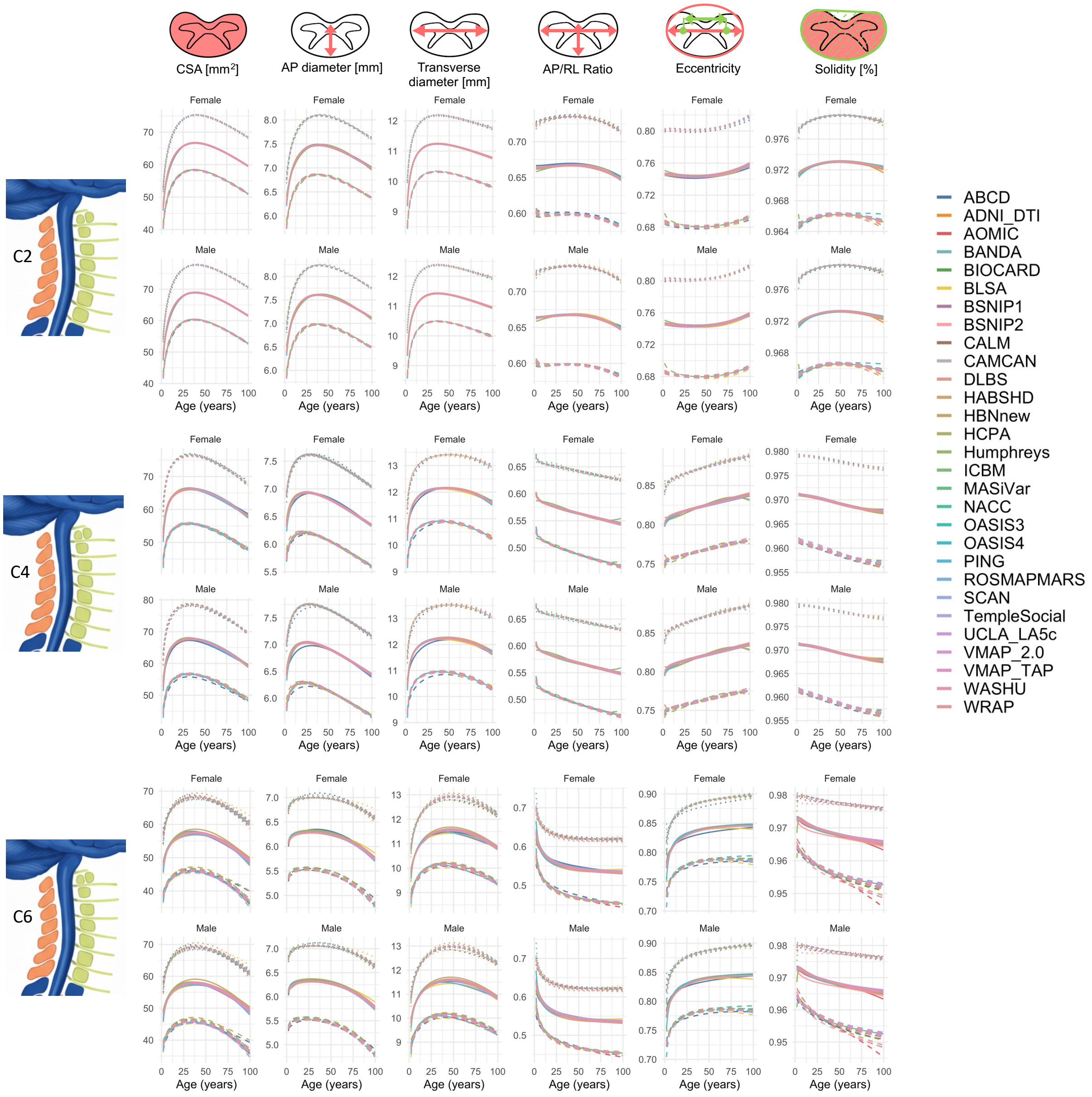


***Supplementary Figure 8**. Leave-one-out site analysis to assess robustness of models to specific sites. Visually, centiles scores remain similar, with more site-driven variation at more caudal levels. Left-out datasets are shown as different colors.*

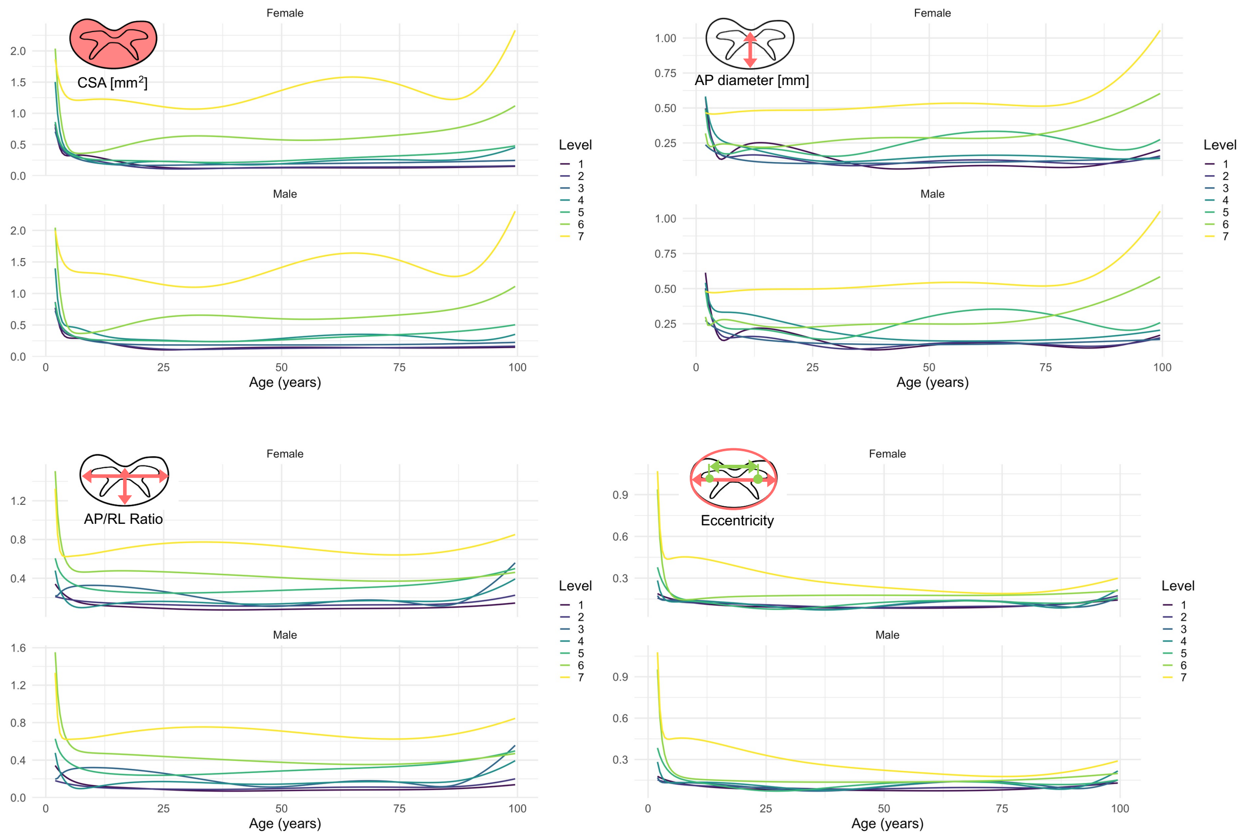


***Supplementary Figure 9.*** *Leave-one-out site analysis to assess robustness of models to specific sites. Percent coefficient of variation across leave-on-out runs, for all ages. CV is typically below 1% (with exception of C7), and often below 0.3% for most features and ages, suggesting a low sensitivity to study composition.*

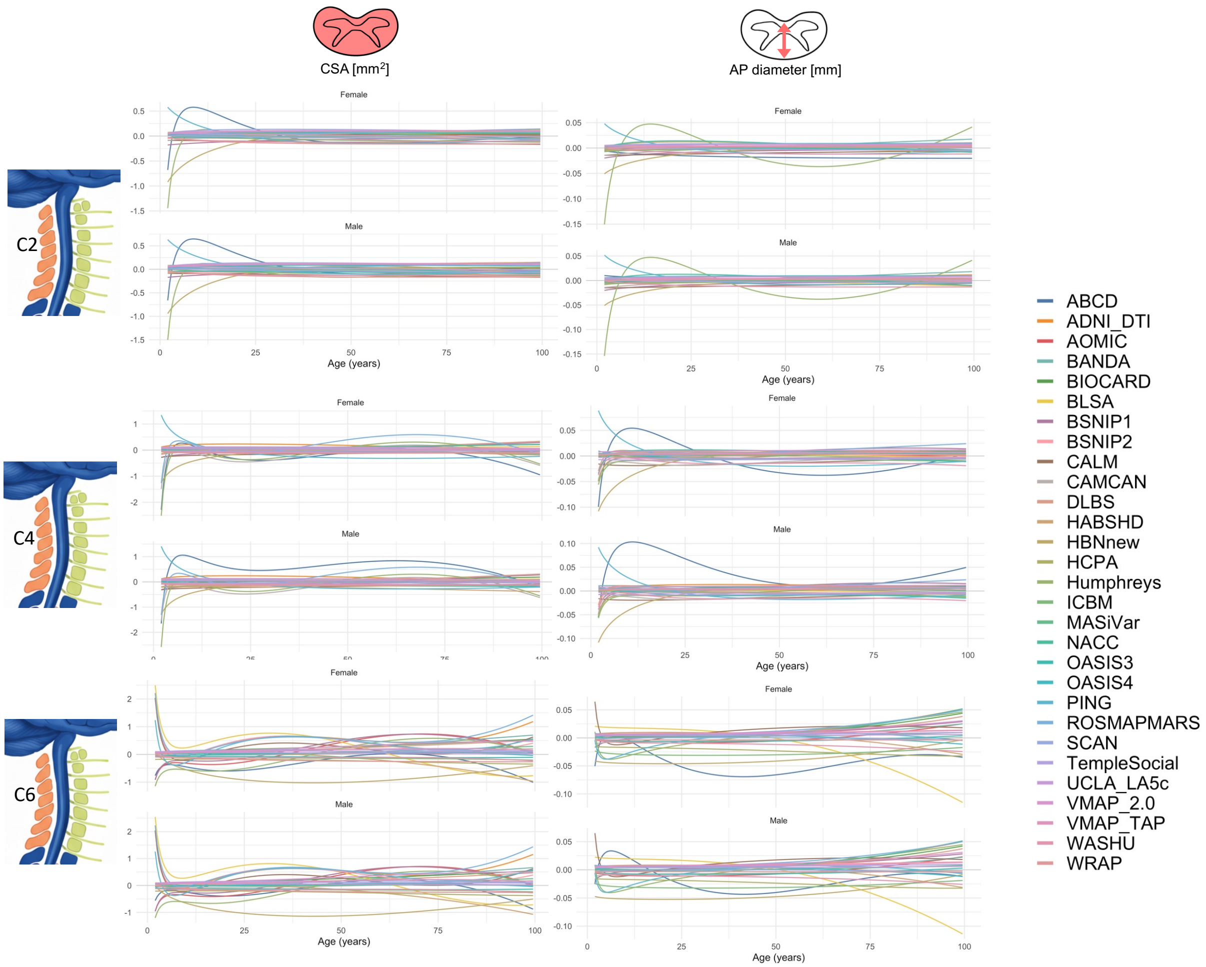


***Supplementary Figure 10.*** *Leave-one-out site analysis to assess robustness of models to specific sites. Plots show the magnitude of influences comparing each left-out-site to the baseline when all datasets are included. CSA, for example, shows <0.5mm^3 differences due to study composition.*

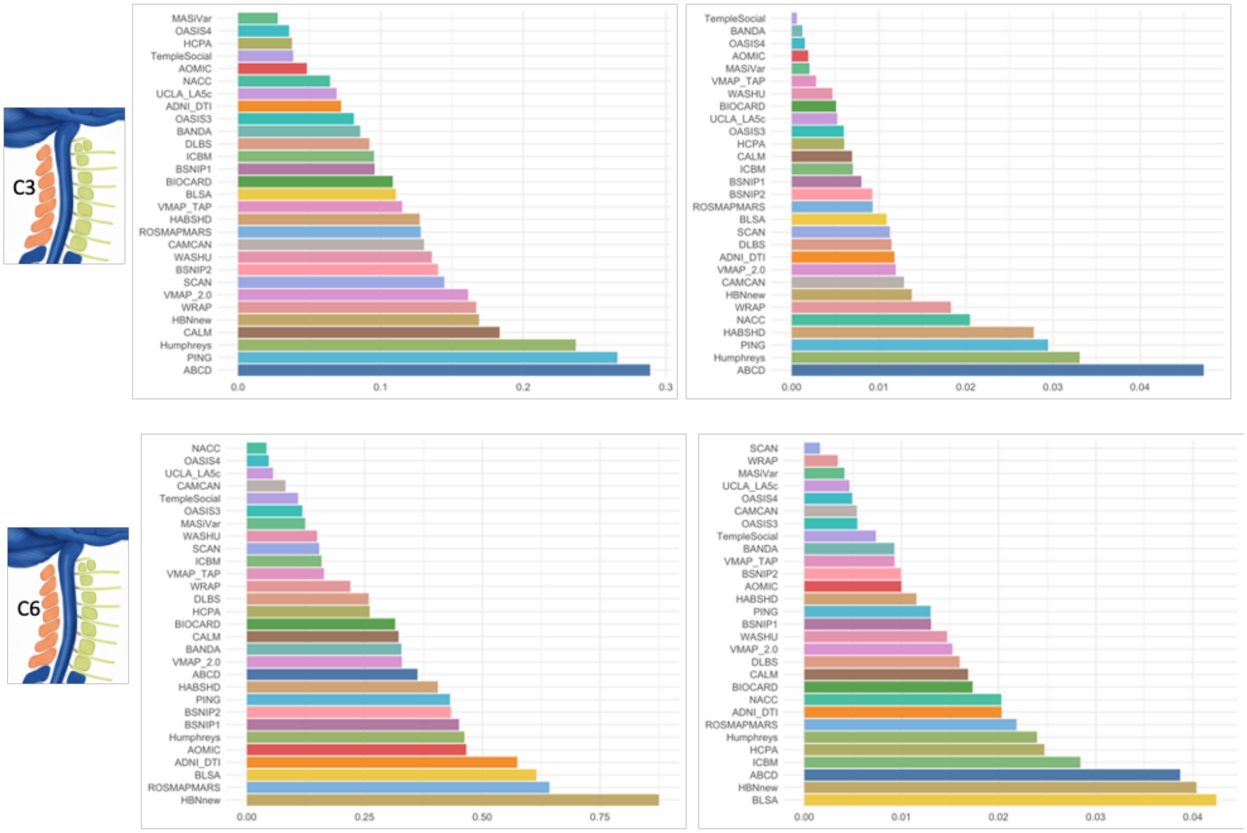


***Supplementary Figure 11.*** *Leave-one-out site analysis to assess robustness of models to specific sites. Plots show the magnitude of influences comparing each left-out-site to the baseline when all datasets are included – taking the absolute value of the magnitude of differences from baseline. Intuitively, sites with larger sample sizes (ABCD, BLSA) have greater overall effects, while sites with lower ages (Humphreys) have greater influence at that lifespan stage.*
